# ATR and PKMYT1 inhibition re-sensitize a subset of TNBC patient-derived models to carboplatin inducing mitotic catastrophe

**DOI:** 10.1101/2025.01.07.629892

**Authors:** Juliet Guay, Hellen Kuasne, Catherine Chabot, Kathryn Bozek, Yasamin Majedi, Marguerite Buchanan, Adriana Aguilar-Mahecha, Eric Bareke, Tim Kong, Kangning Yang, Oluwadara Elebute, Ruining Guo, Anie Monast, Geneviève Morin, Sidong Huang, Morag Park, Mark Basik

**Author notes:** **Corresponding author**: Mark Basik, Lady Davis Institute for Medical Research, Jewish General Hospital, 3755 Côte Ste-Catherine Road, Montreal, Qc, H3T 1E2.

## Abstract

Triple negative breast cancer (TNBC) is associated with poor prognosis and is mainly treated with chemotherapy-based regimens, often including carboplatin. Resistance to carboplatin is a common clinical issue that is either initially present or develops with treatment. Overcoming this resistance is a significant clinical challenge, which highlights the need for novel therapeutic strategies. We used a pooled shRNA screening approach with a chemoresistant TNBC patient-derived xenograft (PDX) cell (PDXC) line to identify targets whose knockdown would enhance the efficacy of carboplatin. This screening led to the identification of the ATR (ataxia telangiectasia and Rad3-related) gene as a key therapeutic vulnerability. Inhibiting ATR with BAY1895344 or AZD6738 re-sensitized carboplatin-resistant PDXCs and PDXs to carboplatin, resulting in an increase in DNA damage, and apoptosis. ATR inhibition disrupts the dependence of carboplatin-resistant cells on the S and G_2_/M checkpoints for DNA repair, leading to mitotic catastrophe. We further found that the addition of ATR inhibitors to carboplatin reversed a FOXM1-targeted gene program enabling premature passage into mitosis. Moreover, targeting PKMYT1, a regulator of cyclin-dependent kinase 1 (CDK1) controlling the G_2_/M checkpoint, through knockdown or with the novel PKMYT1 inhibitor RP-6306, also enhanced carboplatin efficacy in our TNBC PDXC. Molecular factors associated with response to the ATR inhibitor/carboplatin combination included low RNA levels of PKMYT1. These results underscore the pivotal roles of ATR and PKMYT1 in mediating resistance to carboplatin in TNBC and support targeting these pathways to overcome carboplatin resistance in this disease.

## Background

Triple negative breast cancers (TNBC) do not express estrogen receptor (ER), progesterone receptor (PR) nor have overexpression or amplification of the epidermal growth factor 2 receptor (HER2) and are associated with a poor prognosis^1–3^. Despite recent advances in TNBC treatment that have led to the recent approvals of antibody-drug conjugates like Sacituzumab govitecan, PARP inhibitors like olaparib and talazoparib, and immunotherapies like atezolizumab and pembrolizumab, TNBC treatment still relies heavily on chemotherapy, including in the neoadjuvant setting in early TNBC^4^. A key component of chemotherapy in TNBC is carboplatin, as it was shown to improved pathological complete response (pCR) in combination with anthracyclines and taxanes^5^. Nonetheless, at least 35% of early TNBCs are resistant to carboplatin-containing neoadjuvant therapy and metastatic TNBC patients inevitably develop resistance to carboplatin. Uncovering ways to overcome carboplatin resistance in TNBC remains a major unmet need in this disease.

Carboplatin is a platinum based chemotherapeutic drug, an alkylating agent that works primarily by binding to DNA inducing DNA crosslinks, thereby interfering with DNA synthesis and inducing replicative stress, ultimately leading to cell death. Several studies have uncovered different mechanisms involved in carboplatin resistance including decreased drug uptake, increased drug detoxification, increased DNA repair processes and repression of apoptotic signalling among others^6^. In TNBCs, carboplatin resistance has been associated with deregulation of the mitotic checkpoint^7^. However, none of these mechanisms have led to successful therapeutic treatments to overcome carboplatin resistance in the clinic.

Deficiencies in DNA repair pathways, including homologous recombination repair, particularly due to loss-of-function mutations in the BRCA1 and BRCA2 genes, are linked to increased sensitivity of TNBCs to carboplatin^8^. This suggests that the combination of carboplatin with targeted therapies that disrupt the DNA damage response (DDR), may be able to overcome resistance to carboplatin. ATR is a serine/threonine kinase that plays a pivotal role in the DDR and replicative stress. ATR regulates the cell cycle in response to DNA damage via its main effector CHK1 in S phase and in G2/M^9,10^ leading to cell cycle arrest and facilitating DNA repair. In the presence of stalled replication forks, ATR mediates the activation of the mitotic inhibitor WEE1, through CHK1 activation, which results in the inactivation of CDK1 preventing entry into M phase.

ATR inhibition can significantly sensitize cancer cells to DNA damaging agents such as chemotherapy and radiation as shown in extensive pre-clinical and clinical studies^11–13^. Defects in DNA damage response mechanisms, such as ATM mutations affecting homologous recombination, dysfunctional TP53 and elevated levels of replication stress correlate with sensitivity to ATR inhibitors^14,15^. Several ATR inhibitors are in pre-clinical development and are being tested in Phase I and Phase II clinical trials as single agents or in combination with DNA damaging agents or with immune checkpoint inhibitors^14^. Notably, berzosertib (M6620/VX-970/VE-822)^16^ and ceralasertib (AZD6738)^17^ have been tested in combination with platinum agents and have demonstrated antitumor activity in advanced solid tumors, Elimusertib (BAY1895344) is currently being tested in combination with cisplatin in urothelial cancer and results have not been reported yet. Only a few studies have tested ATR inhibitors in the context of platinum resistance. Lung cancer cells deficient in TP53 and ERCC1 treated with an ATR inhibitor were resensitized to cisplatin by inducing replication catastrophe^18^. In ovarian cancer cells ATR inhibition increased the sensitivity to cisplatin though deregulation of its downstream kinases CHK1 and WEE1 and through inhibition of homologous recombination^19^.

PKMYT1 is a crucial cell cycle modulator as it phosphorylates and inhibits CDK1, preventing unscheduled mitotic entry^20–22^. In CCNE1 amplified tumors, including TNBCs, a novel PKMYT1 inhibitor, RP-6306, was found to be synthetic lethal^23^. PKMYT1 inhibition in combination with gemcitabine was shown to reverse resistance to endocrine therapy and palbociclib in ER+ breast cancers^24^. Moreover, PKMYT1 inhibition can synergize with the alkylating agent temozolomide in glioblastoma^25^. A phase I clinical trial testing the PKMYT 1 inhibitor, RP-6306 with carboplatin and paclitaxel in TP53 mutated ovarian and uterine cancer (NCT06107868) is currently ongoing. Interestingly, the synthetic lethality of RP-6306 is also being evaluated in clinical trials with ATR inhibitors in solid tumors.

In the present study, we established patient-derived xenograft cell lines (PDXCs) from chemoresistant TNBCs and performed a pooled shRNA based synthetic lethal screen that led to the identification of ATR as the top synthetic lethal target that resensitized TNBCs to carboplatin. We demonstrate that carboplatin synergizes with the ATR inhibitor BAY1895344 in vitro and that ATR inhibition can resensitize TNBC PDXs to carboplatin and also prevent tumor relapse. We also found PKMYT1 inhibition can overcome carboplatin resistance in TNBCs. The ATRi/carboplatin and PKMYT1i/carboplatin combinations led to accumulation of cells in G2/M and mitotic catastrophe. Together, these results highlight the vulnerability of TNBCs to cell cycle regulators when combined with the DNA-damaging agent carboplatin, supporting further clinical investigation of these combinations in the context of carboplatin resistant TNBC patients.

## Results

### An shRNA screen in carboplatin-resistant TNBC PDX cells identifies ATR as a synthetically lethal gene with carboplatin

To identify molecular vulnerabilities that can be therapeutically exploited to overcome resistance to carboplatin in TNBC, we performed a high throughput shRNA screen using a carboplatin resistant patient-derived cell line model (T-786). The PDX model was generated from a patient who was resistant to carboplatin in both the neoadjuvant and metastatic settings. Tumor cells from the PDX were collected at an early passage to establish PDXCs using a conditionally reprogrammed cell protocol^26^. PDXCs contain the same genomic changes as in both the original tumor and the PDXs from which they were derived (**Supplementary Figure S1A**). *In vitro* resistance was confirmed by cell viability assays (IC_50_ = 52μM) (**Supplementary Figures S2A and E**). T-786 PDXCs were infected with pooled lentiviral shRNA libraries against FDA-approved drug targets^27^ and the human kinome^28^ (∼ 1700 druggable genes). Cells were cultured for 14 days with IC_25_ of carboplatin or vehicle and the relative abundance of shRNA vectors identified by next generation sequencing was quantified followed by model-based analysis of genome-wide CRISPR-Cas9 knockout (MAGeCK) (**Figure 1A**). Ataxia telangiectasia and Rad3-related (ATR) gene was the top candidate whose knockdown was synthetically lethal with carboplatin (**Figure 1B and Supplementary file 1**). The other four top targets identified included CDK2, PLAU, BRAF and ABCC4. Of these, CDK2 and BRAF are druggable but clearly less potent in resensitizing to carboplatin than ATR. Moreover, BRAF is not mutated in this model.

**Figure 1:**
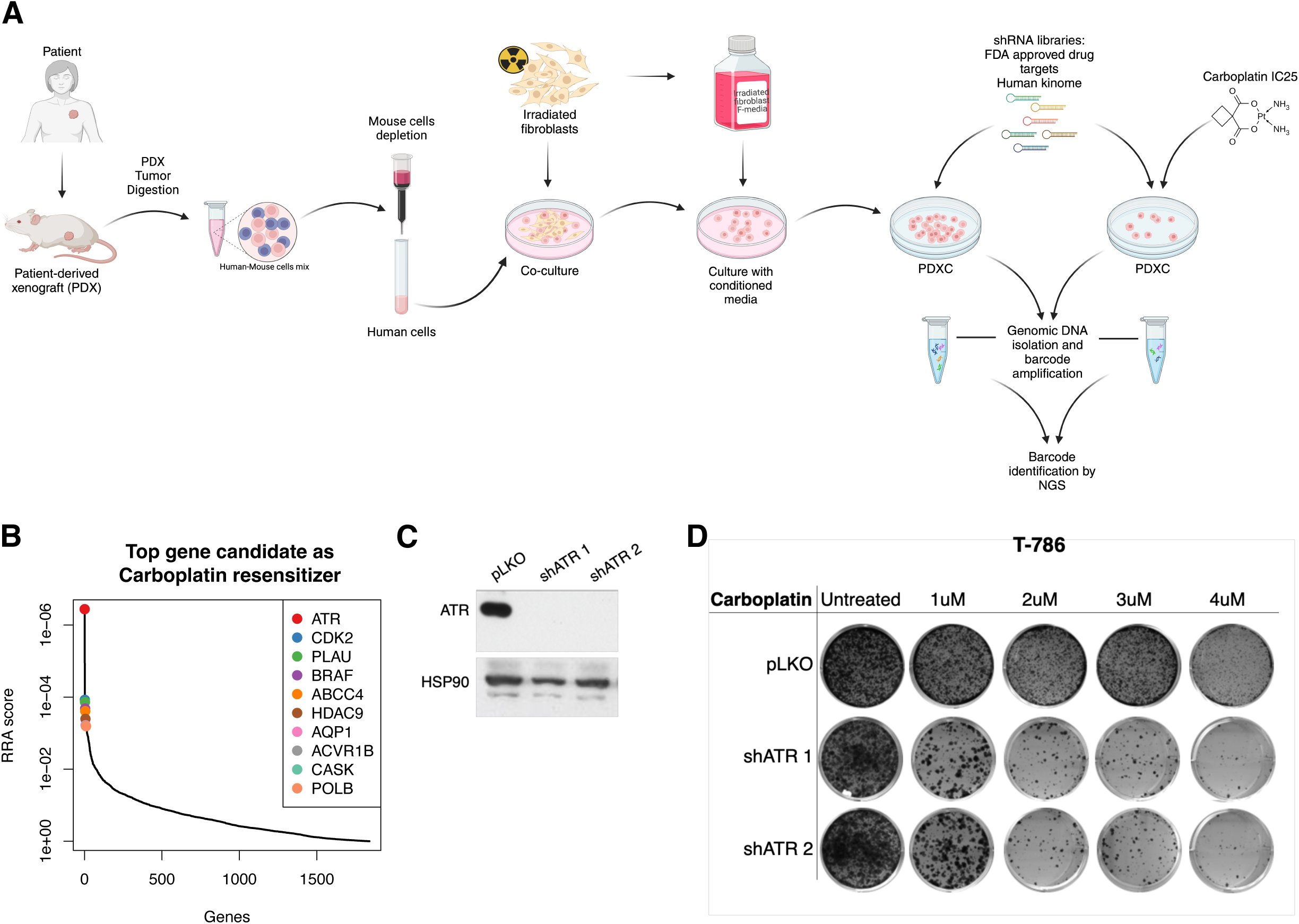
An shRNA screen reveals ATR as the top carboplatin sensitizer gene in a carboplatin-resistant TNBC patient-derived xenograft cell line (T786 PDXC). **A**. Schematic representation of the patient-derived xenograft (PDX) and patient-derived xenograft cell lines (PDXCs) generation and the shRNA screen conducted to identify genes synthetically lethal with carboplatin. Created in BioRender (agreement number: XD27CR1XAU). **B**. Plot of the top 10 carboplatin sensitizer genes rank-ordered by robust rank aggregation (RRA) scores calculated by MAGeCK, where a smaller RRA score indicates more essentiality. ATR gene appeared as the top carboplatin sensitizer gene in PDXC T-786. **C**. Immunoblot analysis of ATR in T-786 transduced with two independent ATR shRNAs. **D**. Clonogenic assay with the two ATR shRNAs-transduced T-786 PDXCs. Cells were treated for 14 days with the indicated concentrations of carboplatin.

### ATR inhibition re-sensitizes chemoresistant TNBC models to carboplatin

To validate these results, we silenced ATR using two shRNAs in the same model. While ATR silencing alone had little impact on colony formation of T-786 cells, it strongly suppressed cell growth when combined with carboplatin (**Figure 1D**). Consistent with our RNAi results, the combination of the ATR inhibitor BAY1895344 (Elimusertib) with carboplatin significantly reduced colony formation (**Figure 2A**). Moreover, BAY1895344 reversed carboplatin resistance in four PDXC models in cell viability assays (**Figure 2B**). We also tested a second ATR inhibitor, AZD6738 (Ceralasertib), which was less potent than BAY1895344, with an IC_50_ > 9-fold higher than BAY1895344 in T-786 (**Supplementary Figure S2E**). BAY1895344 yielded a stronger synergistic combination index than AZD6738 in all PDXCs tested in combination with carboplatin (**Figure 2C and Supplementary Figure S3**).

**Figure 2:**
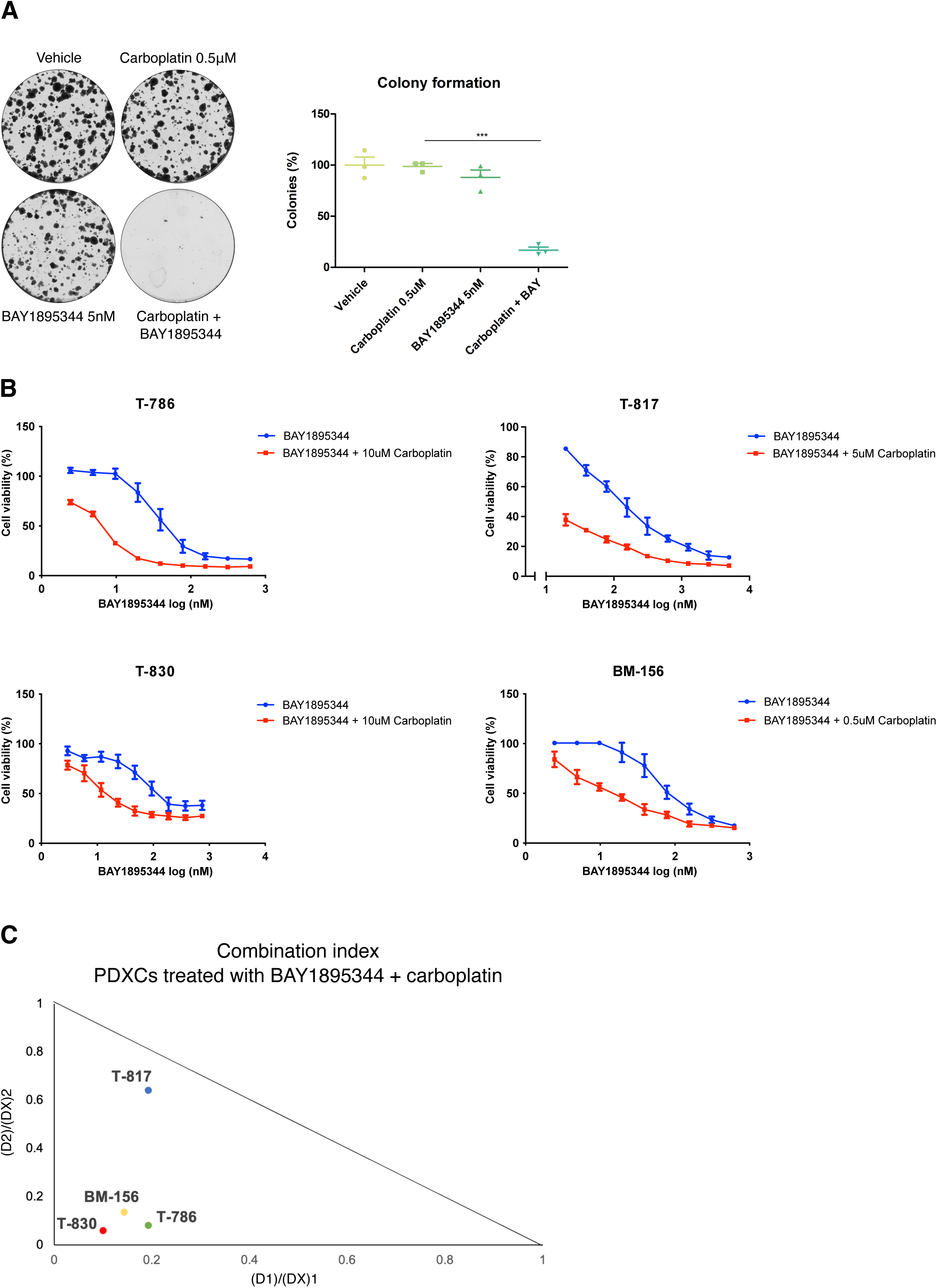
Pharmacological inhibition of ATR synergizes with carboplatin in TNBC PDXCs. **A**. Representative images of clonogenic assay of PDXC T-786 treated with vehicle, 5nM BAY1895344, 0.5μM carboplatin and 5nM BAY1895344 + 0.5 μM carboplatin (left), and the quantification (right) for 14 days. Significance assessed by the two-sided unpaired nonparametric Student’s t test. Mean ± SEM, n=3, ***P < 0.001. **B**. Cell viability (%) of PDXCs T-786, T-817, T-830 and BM-156 measured by AlamarBlue assay. Carboplatin in concentrations averaging between the IC_10_ and IC_25_ concentrations for each cell model was added to a gradient concentration of BAY1895344 (n=3). **C**. Isobolograms representing the combination indexes of each TNBC PDXCs treated with the combination of BAY1895344 and carboplatin.

We next performed *in vivo* tests of the combination of carboplatin with both ATR inhibitors in two matching PDXs, from which the PDXCs were originally derived. We first determined that 20mg/kg of carboplatin with 40mg/kg of BAY1895344, both given once weekly was well tolerated in NSG mice (**Supplementary Figure S4**). BAY1895344 significantly enhanced carboplatin efficacy resulting in a greater tumor growth inhibition (TGI) (TGI = 40%) than the AZD6738 combination (TGI = 14%) in PDX T-786 (**Figure 3A**). Moreover, the combination of either ATR inhibitors with carboplatin significantly improved survival in this model (**Figure 3B**). In model BM-156, Carboplatin combined with BAY1895344 or AZD6738 resulted in TGI of 40% and 65% respectively (**Figure 3C**) and the combination with AZD6738 greatly extended mice survival (**Figure 3D**). Collectively these results support that ATR inhibition and carboplatin treatment offer a promising therapeutic avenue to treat carboplatin resistant TNBC.

**Figure 3:**
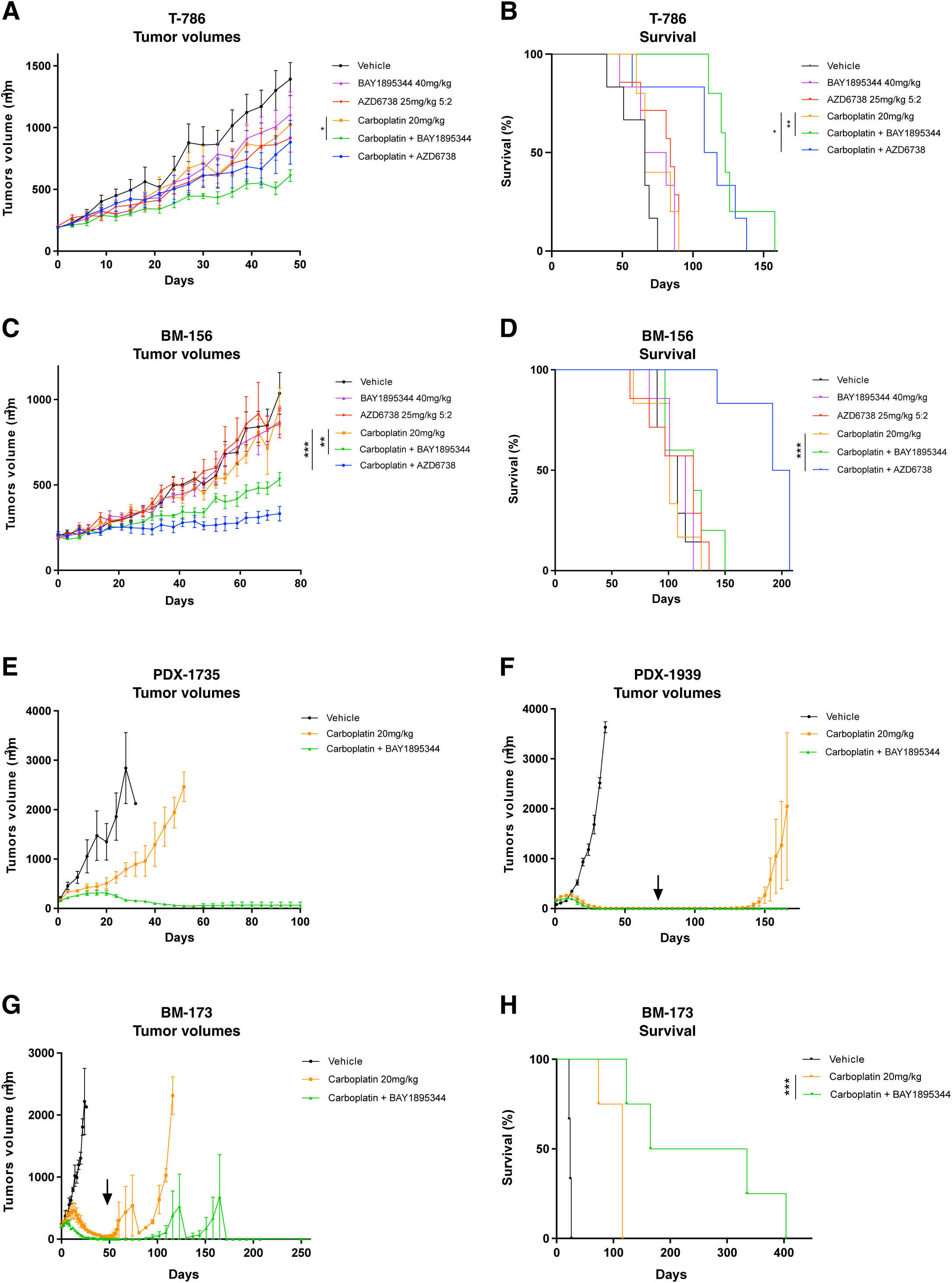
ATR inhibition resensitizes chemoresistant TNBC PDXs and TNBC PDXs that acquire resistance to carboplatin. **A**. Tumor growth and **B.** Survival in response to vehicle (n=5), carboplatin 20mg/kg once weekly (n=7), BAY1895344 40mg/kg once weekly (n=6), AZD6738 25mg/kg 5 days on 2 days off (n=6), carboplatin 20mg/kg once weekly + BAY1895344 40mg/kg once weekly (n=6), or carboplatin 20mg/kg once weekly + AZD6738 25mg/kg 5 days on 2 days off (n=6) in PDX T-786. **C**. Tumor growth and **D.** Survival in response to vehicle (n=5), carboplatin 20mg/kg once weekly (n=6), BAY1895344 40mg/kg once weekly (n=6), AZD6738 25mg/kg 5 days on 2 days off (n=6), carboplatin 20mg/kg once weekly + BAY1895344 40mg/kg once weekly (n=6), or carboplatin 20mg/kg once weekly + AZD6738 25mg/kg 5 days on 2 days off (n=6) in PDX BM-156 **E.** Tumor growth in response to vehicle (n=3), carboplatin 20mg/kg once weekly (n=3), or carboplatin 20mg/kg once weekly + BAY1895344 40mg/kg once weekly (n=3) in PDX-1735. **F.** Tumor growth in response to treatments (see **E**) in PDX-1939 (n=3 per group). **G.** Tumor growth in response to treatments (see **E**) in PDX BM-173 (n=4 per group). **H.** survival in response to treatments (see **G**) in PDX BM-173 (n=4 per group). Mean ± SD, ns = nonsignificant; *P < 0.05, **P < 0.01, ***P < 0.001. Significance assessed by unpaired nonparametric Mann-Whitney of tumor size at the last day where the study contains all individuals (tumor growth) or by Mantel-Cox test for survival. Arrows represent where treatments stopped (**F** and **G**).

We screened an independent series of 12 TNBC PDX models for efficacy of the ATRi/carboplatin combination compared to carboplatin alone using small n=3 experiments (**Supplementary Figure S5 and Figure 3E-H**). Of these, 9 showed varying degrees of resistance to carboplatin while 5 were sensitive (**Supplementary table S5**). The addition of BAY1895344 to carboplatin resulted in tumor regression in PDX-1735 (**Figure 3E**). Furthermore, we observed that acquired resistance to carboplatin was significantly delayed if not prevented by the addition of BAY1895344 in 2 carboplatin sensitive PDX models, PDX-1939 and PDX BM-173, (**Figure 3F-G**). Notably PDX-1939 tumors never regrew even after treatment was stopped, while tumors regrew in the carboplatin arm. BM-173 mice on the combination treatment had four times longer average survival compared to carboplatin treated mice (**Figure 3H**). Together, these results confirm that combining ATR inhibition to carboplatin is a therapeutic strategy that can prevent tumor relapse in a subset of TNBC.

### Pharmacological inhibition of ATR leads to a decrease in its protein expression

We next studied the effects of ATR inhibition on ATR signaling, DNA damage and the cell cycle. ATR autophosphorylation was increased by carboplatin and completely abolished when BAY1895344 was added and decreased when AZD6738 was added (**Figure 4A**). Once activated, ATR phosphorylates its primary effector CHK1 at Ser317 and Ser345^29,30,31^. In the T786 PDXC model, carboplatin alone resulted in CHK1 phosphorylation at Ser345 (**Figure 4A**), and the addition of ATR inhibitor resulted in a marked decrease in this CHK1 activation (**Figure 4A**). Interestingly, the combination of carboplatin with Rabusertib, a CHK1 inhibitor, was not as synergistic as the combination with ATR inhibitors (Chou-Talaly combination index 0.88), suggesting that CHK1 may not be as critical as ATR for survival during carboplatin treatment of these resistant cells (**Supplementary Figure S7**). Indeed, CHK1 did not emerge as a candidate in the original shRNA screen. These findings suggest that ATR signaling independent of CHK1 activation may be at least partly responsible for resistance to carboplatin in these cells.

**Figure 4:**
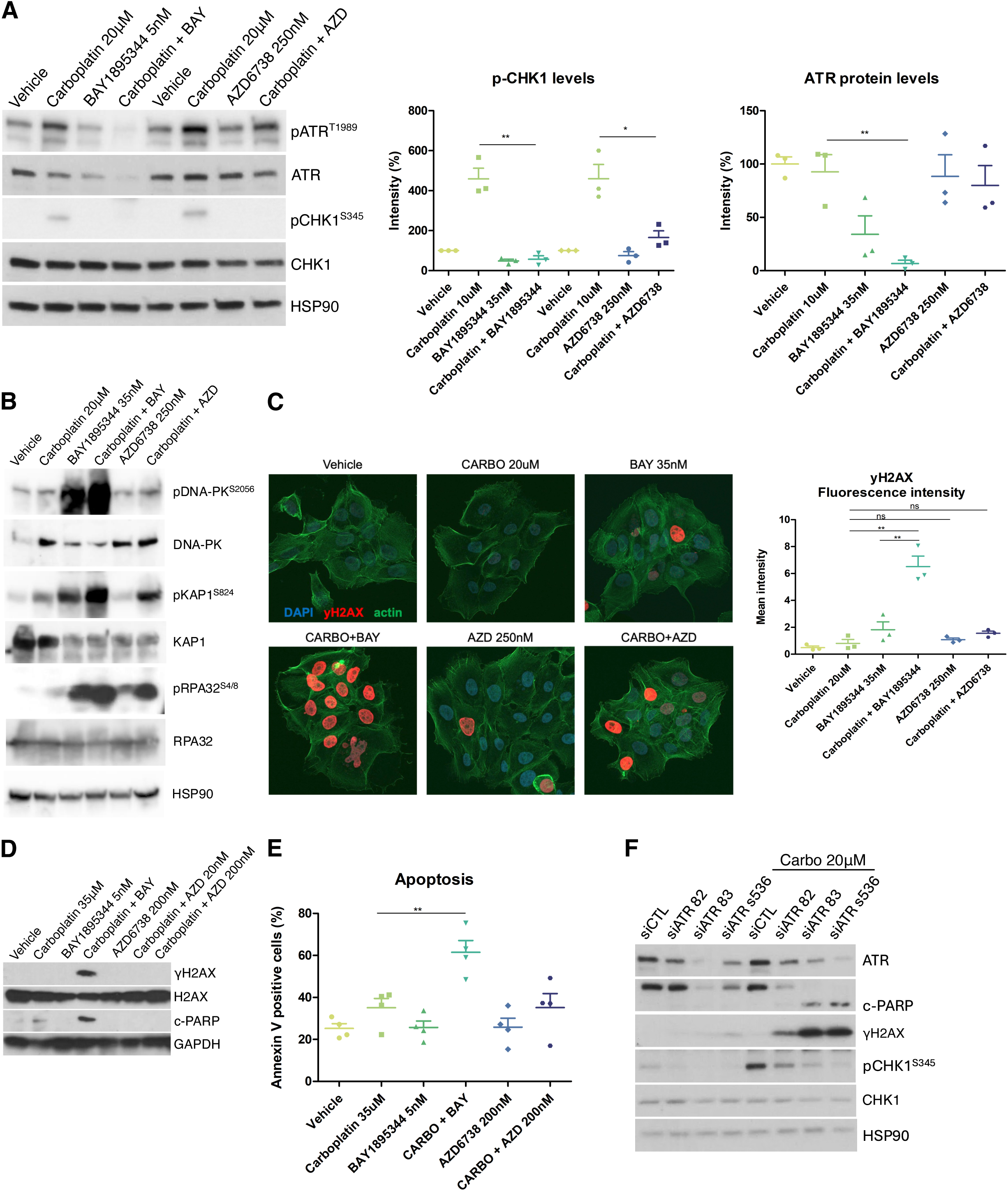
ATR inhibition and knockdown in Carboplatin-treated a TNBC PDXC induces DNA damage and apoptosis. **A**. Immunoblot analysis showing inhibition of ATR activation (p-ATR T1989), and downstream signaling (p-Chk1 S345), in response to 48h of the indicated treatments in PDXC T-786. Quantification of p-CHK1 (S345) and ATR total levels are shown on the right. Significance assessed by the two-sided unpaired nonparametric Student’s t test. Mean ± SD, n = 3, ns, not significant; *P < 0.05, **P < 0.01. **B**. Immunoblot analysis showing stronger induction of the DNA-damage response (p-DNA-PK S2056 and p-KAP1 S824) and replicative stress markers (p-RPA32 S4/8) when ATR inhibitors are combined with carboplatin compared to carboplatin alone in PDXC T-786. **C**. Representative images (left) of immunofluorescence of γH2AX levels (red) acquired by confocal microscopy and γH2AX levels quantification (right). Significance assessed by the two-sided unpaired nonparametric Student’s t test. Mean ± SD, n = 3, ns, not significant; **P < 0.01. Arrow shows the pan-nuclear γH2AX phenotype. **D**. Immunoblot analysis of DNA damage (γH2AX) and apoptotic marker (c-PARP) in response to the indicated treatments in PDXC T-786. **E**. Annexin V levels measured by flow cytometry in response to the indicated treatments in PDXC T-786. Significance assessed by the two-sided unpaired nonparametric Student’s t test. Mean ± SD, n = 3, ns, not significant; **P < 0.01. **F**. Immunoblot analysis showing ATR knockdown in PDXC T-786 treated with three independent siRNAs and the effect on downstream signaling (p-CHK1 S345), DNA damage (γH2AX) and apoptosis (c-PARP).

Although carboplatin resulted in autophosphorylation of ATM, this activation was not abrogated when combined with ATR inhibitors, confirming the specificity of these ATR inhibitors (**Supplementary Figure 8A**). We observed that ATR protein levels decreased with BAY1895344 alone and further when combined with carboplatin (**Figure 4A**). This ATR protein loss was also observed in other TNBC PDXCs (**Supplementary Figure S8B**). We did not find any involvement of transcriptional regulation or proteasomal degradation to explain this decrease in protein levels (**Supplementary Figure S8C-E**). A cellular fractionation assay did not reveal ATR sequestration on the chromatin in response to the combination (**Supplementary Figure 8F**). While the mechanism underlying suppression of ATR protein expression by BAY1895344 remains unclear, this unexpected effect may contribute to the effect of the BAY1895344 and carboplatin combination *in vitro*.

### Carboplatin in combination with ATR inhibitors leads to DNA damage, replicative stress, mitotic catastrophe and cell death

We further studied the molecular effects of carboplatin-ATR inhibitor combination treatment in PDXCs. Carboplatin treatment alone resulted in a slight induction of p-RPA32, a marker of replicative stress ^32–34^ (**Figure 4B**), but not of γH2AX, a marker for double strand breaks^35^ in PDXC T-786 (**Figure 4C**). Treatment with BAY1895344 alone resulted no significant induction of γH2AX but induction of p-DNA-PK, p-KAP1, a substrate of ATM induced by double strand breaks^36^ and p-RPA32 which was exacerbated by the addition of carboplatin. The combination of carboplatin with BAY1895344 resulted in marked increases in γH2AX and p-RPA32 (**Figure 4C-D**). Interestingly, the pattern of γH2AX fluorescence was uniform throughout the nucleus or pan-nuclear, suggestive of excessive replicative stress, mitotic catastrophe, and cell death^37^ (**Figure 4C**). After 3 days of treatment, these cells started to undergo apoptosis as evidenced by increased levels of cleaved-PARP and annexin V (**Figure 4D-E**). Higher doses of AZD6738 could initiate a similar response (**Supplementary figure S9)**, pointing out to a similar mechanism of action than BAY1895344 but with a lower potency. Genetic inhibition of ATR using three different siRNA also showed DNA damage and apoptosis as indicated by γH2AX and cleaved-PARP in carboplatin treated cells (**Figure 4F**). Altogether, these results show that pharmacologic and genetic ATR inhibition combined with carboplatin induces DNA damage and increases replicative stress leading to apoptosis, demonstrating that ATR signaling is essential for the survival of carboplatin treated T-786 cells.

It is well established that ATR activation can trigger cell cycle arrest at the S phase by CHK1 dependent inhibition of CDC25A phosphatase activity on CDK2^9,10,38,39^. Thus, ATR inhibition should reverse cell cycle arrest at the S phase. More recent evidence shows that ATR controls the S/G_2_ checkpoint by delaying FOXM1 phosphorylation until G_2_, thus inhibiting the transcription of mitotic gene networks^9^. In PDXCs T-786, ATR inhibition alone did not result in any appreciable change in cell cycle dynamics. On the other hand, carboplatin alone resulted in an increase in the number of cells in S and G_2_/M phases (**Figure 5A**), consistent with an increased time for repair of carboplatin induced DNA damage before mitosis. When ATR was inhibited, the addition of carboplatin resulted in a marked accumulation of cells in G_2_/M, suggesting that ATR prevents S/G_2_ progression in carboplatin treated carboplatin-resistant cells. (**Figure 5A**). We observed more mitotic figures in cells treated with the combination, together with micronucleation and nuclear fragmentation (**Figure 5B**), signs of premature mitosis resulting in mitotic catastrophe^40^. Histone H3 was phosphorylated by the combination treatment, which is indicative of mitosis, as was FOXM1, the master regulator of the mitotic gene network expression, indicating its activation (**Figure 5C**).

**Figure 5:**
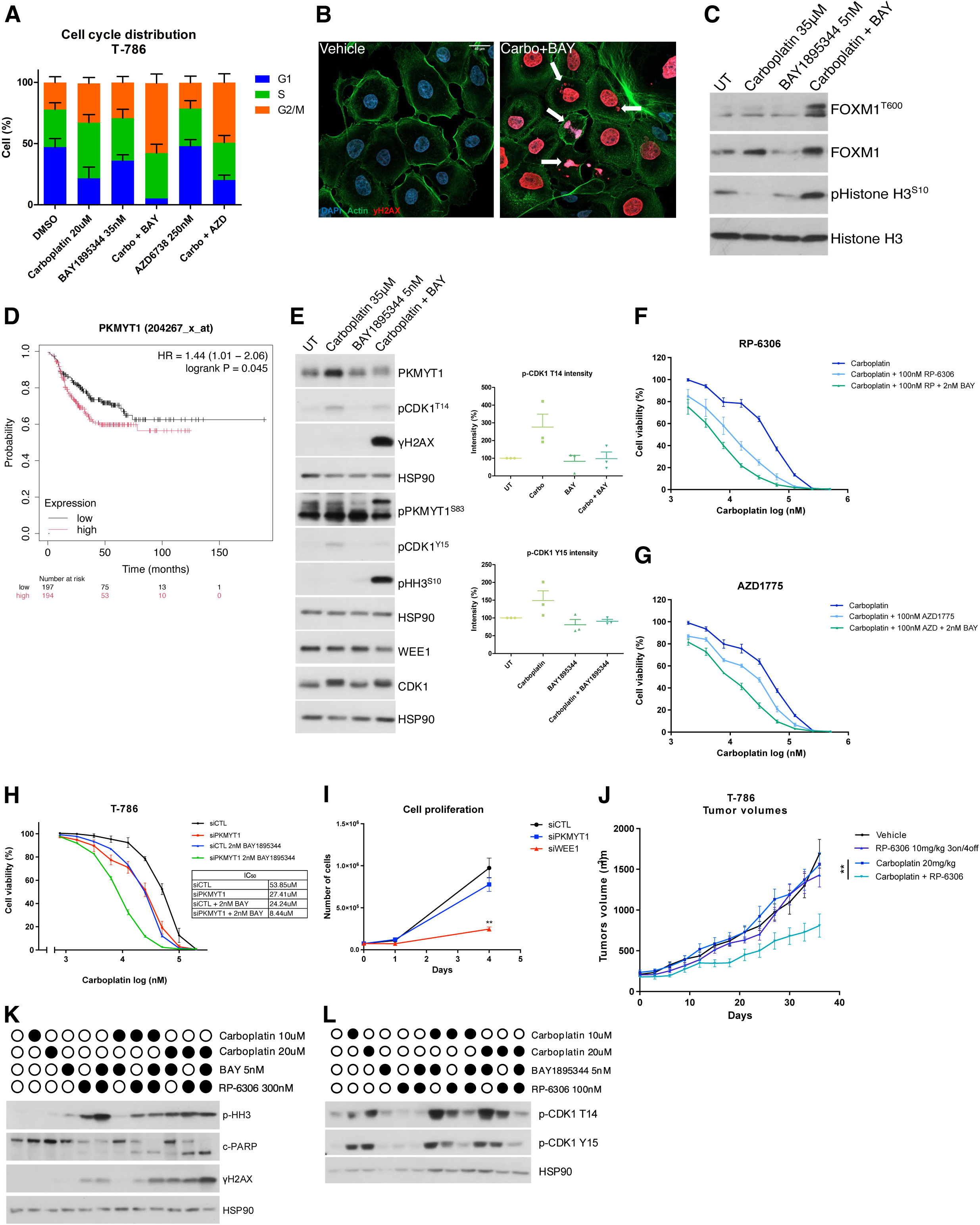
Combination of Carboplatin and ATR inhibitors lead cells to mitotic catastrophe, which involves PKMYT1 inhibition. **A**. Cell cycle distribution of PDXC T-786 quantified by flow cytometry after 24h of treatment with the indicated drug combinations (n=3). **B**. Confocal microscopy images depicting failed mitosis and γH2AX staining (red) in untreated (left) and treated with 20μM carboplatin + 5nM BAY1895344 (right) in PDXC T-786. **C** Immunoblot analysis showing FOXM1 (T600) and histone H3 phosphorylation in PDXC T-786 in response to the indicated treatments. **D**. Immunoblot analysis showing CDK1 regulation, DNA damage (γH2AX) and mitotic entry (pHH3). CDK1 phosphorylation (T14 and Y15) quantification is shown on the right (n=3). **E**. Kaplan-Meier analysis for overall survival of chemotherapy-treated TNBC patients in the KM-plotter cohort with high or low levels of PKMYT1.**F-G**. Cell viability (%) of PDXC T-786 measured by AlamarBlue assay after treatment with the indicated drug combinations (n=3). **H**. Cell viability (%) of PDXC T-786 transfected with a pool of PKMYT1 siRNAs measured by AlamarBlue in response to the indicated treatments. IC_50_ are presented on the right (n=3). **I**. Cell proliferation of PDXC T-786 transfected with PKMYT1 siRNAs (n=3). Number of cells after 24h and 4 days after plating was determined by manual cell counting. **J**. Tumor growth in response to the indicated treatments in PDX T-786. Randomized mice were treated with vehicle, carboplatin 20mg/kg once weekly, RP-6306 10mg/kg twice a day, 3 days on, 4 days off, or the combination of carboplatin and RP-6306. Each treatment arm contains 4 to 6 mice. **P < 0.01, Significance assessed by unpaired nonparametric Mann-Whitney of tumor size. **K**. Immunoblot analysis showing c-PARP, p-HH3 and γH2AX levels in response to 48h of treatments with the indicated drug combinations in PDXC T-786. **L**. Immunoblot analysis showing CDK1 phosphorylation in response to the indicated treatments in PDXC T-786.

### ATR inhibition combined with carboplatin induces the transcription of mitotic processes

To better understand the effect of combining BAY1895344 with carboplatin, we performed RNA sequencing of T-786 cells after 24 hours of treatment. Treatment with Carboplatin alone resulted in the significant differential expression of only 45 genes, of which 39 had their expression suppressed (p adjusted < 0.05) (**Supplementary Figure S13**). Remarkably 37 of these genes are regulated by the FOXM1 transcription factor, according to the ChEA transcription factor targets dataset^41^. A gene ontology analysis (GOA) of the biological processes of these differentially expressed genes revealed an enrichment for upregulated genes involved in the negative regulation of mitosis and for downregulated genes involved in chromosome segregation and mitosis (**Supplementary Figure 14A-B**). These genes included Cyclin B1, Cyclin B2, CDC20, several KIFs, Aurora Kinase A (AURKA) and Polo Like Kinase 1 (PLK1). We then compared the transcriptome of these cells treated with carboplatin alone to that of combination treatment with carboplatin and 5nM BAY1895344 for 24h. DESeq2 analysis revealed that 746 genes were significantly (p adjusted < 0.05) upregulated or downregulated. GO analysis revealed that the combination treatment resulted in a significant enrichment for biological processes related to chromosome segregation division and mitosis for upregulated genes (**Supplementary Figure S14C**), whereas downregulated genes are enriched for the biological processes of DNA replication and cell cycle control (**Supplementary Figure S14D**). Remarkably, all the 39 genes that had been suppressed by carboplatin treatment alone were now significantly upregulated by the addition of an ATR inhibitor, consistent with our observation of FOXM1 phosphorylation with the combination. These findings support that T-786 cells resist carboplatin by inhibiting mitosis and cell division and that this program is tightly regulated by ATR. Moreover, this further supports our findings that mitotic entry is triggered when combining the ATR inhibitor with carboplatin. On the other hand, processes related to DNA replication and cell cycle control were downregulated in response to the combination, suggesting ATR plays an essential role in suppressing replication and slowing the cell cycle to allow for repair of carboplatin-induced DNA damage repair in these carboplatin-resistant cells.

### Inhibition of PKMYT1 re-sensitises cells to carboplatin

Mitotic entry is triggered by CDK1 ^21^ complexing with cyclin B1^42,43^, a process regulated by the inhibitory activity of PKMYT1 and WEE1, through their phosphorylation of Thr14 and Y15 on CDK1. In fact, PKMYT1 can phosphorylate both Thr14 and Y15^44,45^ whereas WEE1 only phosphorylates Y15^46^. PKMYT1 is itself phosphorylated in mitosis which results in its inactivation^45,47,48^. Interestingly, high expression of PKMYT1 is associated with poor prognosis in chemotherapy-treated TNBC patients (**Figure 5D).**

Treatment with carboplatin resulted in increased expression of PKMYT1, but not WEE1, and increased inhibitory phosphorylation of CDK1, suggesting a critical role for PKMYT1 in the regulation of the delay in mitotic entry induced by carboplatin in T-786 (**Figure 5E**). While ATR inhibition alone did not alter the expression or activation of PKMYT1 or CDK1, the combination of BAY1895344 with carboplatin resulted in a marked decrease in PKMYT1 protein levels compared to carboplatin alone. Additionally, the combination led to the phosphorylation of PKMYT1 (**Figure 5E**). PKMYT1 has been shown to be hyperphosphorylated in mitosis^22,47^. The kinases responsible for this phosphorylation are not well documented in humans, but studies in other organisms suggest it could be AKT1^49^, MEK1^50^, p90RSK^51^ and PLK1^48^. Interestingly, PLK1 is one of the genes transcriptionally downregulated by carboplatin treatment, and its expression is upregulated with the addition of BAY1895344 (**Supplementary Figure S13**). We also observed a decrease of the inhibitory T14 and Y15 CDK1 phosphorylation with the combination, consistent with progression of the DNA damaged cells into mitosis, as confirmed by the concomitant increase in γH2AX and pHH3 (**Figure 5D**). All together, these results suggest that PKMYT1 plays a role in the induction of the mitotic delay induced by carboplatin, a function that seems to be decreased when the ATR inhibitor is added.

To better understand the implication of PKMYT1 in carboplatin resistance we tested the novel PKMYT1 inhibitor, RP-6306^52,53^, in combination with carboplatin using a cell viability assay. A clear synergistic effect was observed in PDXC T-786 (CI= 0.52) and the addition of low doses of BAY1895344 further re-sensitized the cells to carboplatin (**Figure 5F**). These results were further validated in the other 3 PDXCs where varying degrees of synergy were observed for the RP-6306 and carboplatin combination: PDXC T-817 (CI=0.53), T-830 (CI=0.85) and BM-156 (CI=0.60) (**Supplementary Figure S10**). In contrast, the WEE1 inhibitor AZD1775 (adavosertib) showed only mild synergy with carboplatin in PDXC T-786, with a combination index close to 0.9 (CI=0.86) (**Figure 5G**). Adavosertib did inhibit WEE1, as shown by the decrease in CDK1 phosphorylation on Y15 when combined with carboplatin (**Supplementary Figure S11C**). Using siRNA silencing of PKMYT1, we found a 2-3-fold increase in sensitivity to carboplatin in PKMYT1 silenced PDXC T-786, and a 6-7-fold increase when a low concentration of BAY1895344 was further added (**Figure 5H**). Similar effects were observed with WEE1 RNAi silencing (**Supplementary figure S11A-B**). However, we noticed a marked effect of the WEE1 knockdown alone on proliferation, contrary to PKMYT1 silencing alone, which showed no effect on proliferation (**Figure 5I**). Given the promising effect of RP-6306 with carboplatin in vitro, we validated this combination in vivo in the matching PDX T-786. RP-6306 had no effect alone but the combination with carboplatin significantly delayed tumor growth (**Figure 5J**).

Since both ATR and PKMYT1 seem to be involved in carboplatin resistance in our models, we investigated whether a triple combination of BAY1895344, RP-6306 and carboplatin could be more efficient at overcoming carboplatin resistance than the double combinations. Indeed, the triple combination led to higher levels of DNA damage and apoptosis than the dual combination of carboplatin with RP-6306 or with BAY1895344 (**Figure 5K**). This correlated with a greater decrease in CDK1 phosphorylation on both T14 and Y15 with the triple combination compared to both double combinations (**Figure 5L**). These findings were further supported by PKMYT1 and ATR double knockdown via siRNAs, which led to increased mitotic catastrophe when combined with carboplatin (**Supplementary figure S12**). Altogether, these results support that PKMYT1 contributes to carboplatin resistance by controlling mitotic entry and that PKMYT1 may be an effective target for carboplatin resistant TNBC cells.

We studied the on-treatment transcriptional changes upon the addition of RP-6306 to carboplatin to compare them with those described above for the BAY1895344/Carboplatin combination. The differentially expressed genes were somewhat different. Treatment with RP-6306 reversed the carboplatin-induced suppression of 26 of the aforementioned 39 genes suppressed by carboplatin, compared to reversal of all of them with ATR inhibition (**Supplementary Table S6**). GOA of the up-regulated genes revealed an enrichment for genes associated with ribosome biogenesis and translation (**Supplementary Figure S13E**), whereas the down-regulated genes were enriched for cell adhesion, cell cycle control and DNA replication (**Supplementary Figure S13F**).

### Transcriptional profiling of responders and non-responders PDXs

To better understand which subset of TNBC might respond to the ATRi/Carboplatin combination and to identify candidate biomarkers predictive of response to the combination, we investigated the genomic and transcriptional differences between models that favorably responded to the combination treatment compared to carboplatin alone. The four models identified as responders were PDX T786, PDX BM-156, PDX-1735, and PDX-1939. PDX-1939 initially showed regression with both carboplatin and the combination but later regrew only with carboplatin and not with the combination, suggesting that the combination prevented the acquisition of carboplatin resistance. It was therefore classified as a responder. In contrast, the six non-responders were PDX-1915, PDX-1924, PDX-1971, PDX-1905, PDX-2076, and PDX-2089 (**Supplementary table S5**). Whole genome sequencing, SNP-CGH and whole exome sequencing was performed on these models. We also focused our attention on 70 genes selected for their role in cell cycle, DNA damage repair and ATR response^54–57^. Interestingly, 1 of these 10 PDX models contained a missense variant of ATM: PDX-1971 (p.Asp2720Asn)^58^ (**Supplementary Figure S6**), but this model was not responsive to the ATRi/Carboplatin combination. A BRCA1 mutation was detected in one of the responders and a BRCA2 mutation in one of the non-responders. As expected, there were few other mutated genes besides TP53. Copy number changes were inferred from whole genome sequencing and SNP-CGH (Cytoscan) and did not reveal obvious differences between responders and non-responders, except for 2 of the 4 responder tumors showing an increase in copy number of the FOXM1 gene. These copy number changes were not focal amplifications however.

We performed RNAseq analysis on all 10 PDX models. Principal component analysis (PCA) revealed distinct clustering of responders and non-responders, highlighting clear transcriptional differences between these groups (**Supplementary Figure S15**). DSeq2 analysis was performed between responders and non-responders. Gene ontology analysis indicated that responders exhibited higher expression of genes associated with cell activation, response to bacterium and angiogenesis (**Figure 6A**) The responders also exhibited lower expression of genes associated with chromosome segregation, organelle fission and nuclear division (**Figure 6B**). We also examined the expression of the selected group of 70 genes (**Supplementary Figure S6**). Notably, 13 of these 70 genes had significantly lower RNA levels in responders relative to non-responders (**Figure 6C**). These included the cell cycle regulator CDC25C, key DNA damage repair genes RAD51, BRCA1, XRCC1, NBN, CHEK2, FANCD2 and ERCC2 as well as PKMYT1. Altogether, these findings suggest that gene expression variation in DNA repair and cell cycle processes may influence the response to the combination of carboplatin and ATR inhibitors. Furthermore, low expression of PKMYT1 may be a potential biomarker for predicting response to this combination therapy, concordant with the increased effects of silencing both PKMYT1 and ATR in reversing carboplatin resistance.

**Figure 6:**
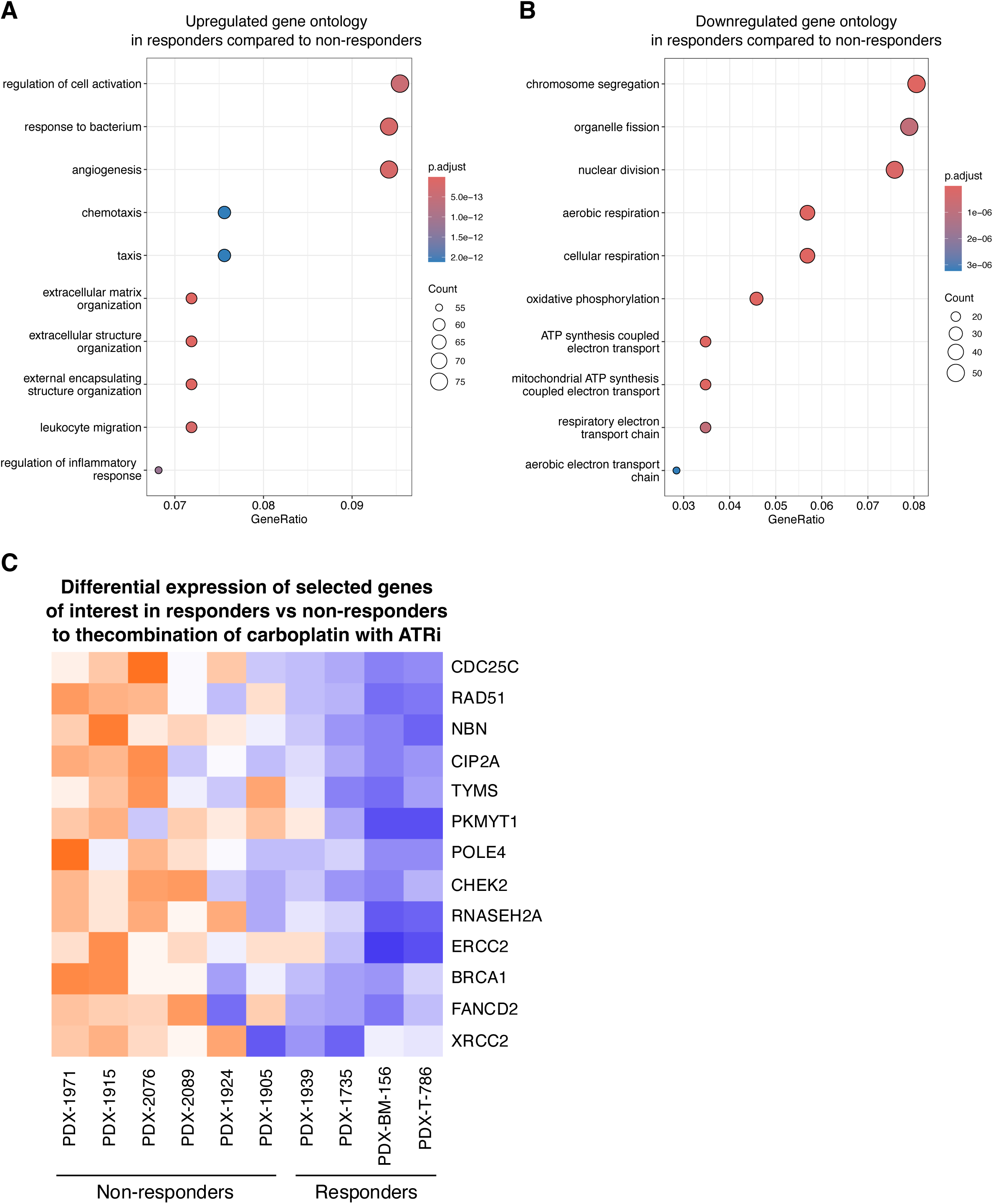
Targeting the cell cycle in carboplatin resistant TNBC **A**. Transcriptomic comparison of responder and non-responder PDXs to the combination of carboplatin + BAY1895344. Dot plot represents the top 10 upregulated and **B**. downregulated gene ontology (GO) biological processes in the PDX that responded to the combination compared to those that did not at basal level, based on differentially expressed genes (DEGs) with *p* adjusted< 0.05. **C**. Heatmap representing the significant transcriptional modulation genes from the gene list used for the model characterization (supplementary figure S6). Represented genes were selected with P < 0.05.

## Discussion

Carboplatin is one of the mainstays of chemotherapy treatment of triple negative breast cancer (TNBC). This DNA damaging agent acts primarily by binding to DNA and producing interstrand and intrastrand cross links^59^. Although carboplatin is highly effective in both early and late TNBC, resistance inevitably develops in the more advanced contexts. Many mechanisms of resistance have been elucidated including alterations in DNA repair mechanisms, which ensure continued viability of the cell^60^. We have generated clinically relevant patient derived models of carboplatin resistance including PDX and PDXCs from drug resistant TNBCs and found through a shRNA screen that ATR was the top gene whose inhibition led to carboplatin re-sensitization. We showed that 2 ATR inhibitors, namely AZD6738 and BAY1895344, can resensitize our PDXCs and PDXs to carboplatin in vitro and in vivo, highlighting the potential for ATR inhibition to overcome carboplatin resistance. Our observation that tumors from PDX-1939 did not regrow following the cessation of treatment is particularly striking, suggesting that the combination therapy may induce long-lasting responses that prevent relapse.

ATR acts to delay cell cycle progression via an intra-S or G_2_/M checkpoint allowing greater time for repair of DNA damage and for resolution of replicative stress^45,70,71^. More recently, ATR has also been implicated in an S/G_2_ checkpoint where it can prevent transcription of mitotic genes via CDK1 and FOXM1, further preventing early mitotic entry^9^ (**Figure 6C**). Our results indicate that ATR inhibition with carboplatin increases DNA damage in carboplatin resistant cells leading to excessive replicative stress and subsequent apoptosis. Our data also revealed that ATR inhibition alone did not significantly alter the cell cycle while the combination with carboplatin led to accumulation of cells in the G2/M phase. This suggests that the inhibition of ATR eliminates a critical checkpoint control that normally prevents cells with damaged DNA from entering mitosis. In fact, we observed an increased presence of mitotic figures, micronucleation, and nuclear fragmentation further supporting the notion that the combination treatment results in premature mitotic entry, leading to mitotic catastrophe. Indeed, PDXC T-786 treated with a combination of ATR inhibition and carboplatin showed high levels of γH2AX concurrently with high levels of p-HH3, suggesting mitotic entry while DNA damage remains unrepaired (**Figure 6D**). Our exploratory analysis of PDXs responding or not responding to the ATR inhibitor/carboplatin combination suggests that the expression levels of specific genes involved in cell cycle regulation and DNA damage repair may serve as valuable biomarkers for predicting the efficacy of this combination.

We found that the combination of carboplatin with ATR inhibition results in FOXM1 phosphorylation, facilitating mitotic entry. RNAseq data from our PDXC models treated with carboplatin or carboplatin and ATR inhibition supports the role of FOXM1, whose target genes involved in chromosome segregation and mitosis are suppressed by carboplatin treatment, delaying mitotic entry. Adding ATR inhibition prevents this suppression, activating mitotic entry. Mitotic entry was further demonstrated by a decrease in the inhibitory phosphorylations of CDK1 (T14 and Y15) in the combination treatment compared to its induction by carboplatin alone. ATR can regulate CDK1 via its effector CHK1 that can inhibit the CDC25 phosphatases, which are responsible for removing these inhibitory phosphorylations^38,39,61^. We found that the knockdown of either kinase responsible for phosphorylating CDK1, WEE1 and PKMYT1, could also resensitize PDXC T-786 to carboplatin, highlighting the pivotal role of the G2/M checkpoint in carboplatin resistance in TNBCs. Contrary to PKMYT1 knockdown, WEE1 knockdown had a drastic effect on cell proliferation. This is in line with PKMYT1 being important for the recovery of the cell cycle after DNA damage while being dispensable for unperturbed cell cycling^22^. The differential effects observed suggest that targeting PKMYT1 could provide a more specific approach to sensitize resistant tumors without substantially impairing normal cell proliferation.

Interestingly, carboplatin treatment increased PKMYT1 expression and high levels of PKMYT1 are associated with poor prognosis in patients with triple-negative breast cancer (TNBC) treated with chemotherapy, suggesting it may play a role in resistance. Moreover, we observed phosphorylation of PKMYT1 in response to the combination of carboplatin with BAY1895344, implying that ATR can regulate this inhibitory phosphorylation (**Figure 6D**).

PKMYT1 has recently gained attention as a promising target, although its therapeutic exploration was previously limited by the lack of specific inhibitors. Using the novel PKMYT1 inhibitor RP-6306^52^, we were able to resensitize carboplatin resistant models highlighting the potential of PKMYT1 inhibition as a therapeutic strategy to enhance the efficacy of carboplatin in drug resistant TNBCs. Although, treatment with RP-6306 was able to partially reverse the carboplatin-induced gene suppression program, other genes were transcriptionally altered in the RP-6306/carboplatin combination, warranting further investigation to understand their role in mediating the synergistic effects with carboplatin. Currently, RP-6306 is being tested in clinical trials in combination with carboplatin and paclitaxel for ovarian and uterine cancers (NCT06107868), and with the ATR inhibitor RP-3500 for advanced solid tumors (NCT04855656). Although our results demonstrate the potential of a triple combination in effectively combating carboplatin resistance in TNBC this triple combination may be overly toxic.

Preclinical investigation of RP-6306 has been in part conducted in the context of CCNE1 amplification or FBXW7 loss^53^ and more recently, in CDK4/6 resistant ER+ breast cancer cells^24^. It is important to note that none of our PDXCs have these biomarkers (**Supplementary Figure S6**), thus further investigation on mechanisms of RP-6306 efficacy in TNBCs is needed.

In conclusion, a subset of carboplatin-resistant TNBCs relies on ATR and PKMYT1 to slow the cell cycle and buy time for the repair machinery to handle DNA damage and replicative stress. Our findings suggest a clinical value for the combination of ATR or PKMYT1 inhibition with carboplatin in carboplatin-resistant TNBCs. Future studies should explore the clinical applicability of these combination therapies in patients with resistant tumors, guided by biomarkers that can predict response and improve personalized treatment strategies in TNBCs.

## Author contributions

JG designed experiments, performed experiments, acquired data, analyzed data, prepared figures, and wrote the manuscript. CC and YM designed experiments, performed experiments, and analyzed data for western blotting and cell cycle analysis. CY designed experiments, performed experiments, and analyzed data for clonogenic assay. HK designed experiments, performed experiments, analyzed data for in vivo studies. KB, AM and MB helped with in vivo studies. RG performed cell viability assay. TK, GM and SH design, conducted and acquired the shRNA screen. AAM and MB design the project, helped with designing the studies, wrote the manuscript and secured funding.

## Materials and methods

### Generation of PDXs and PDXCs

PDXs originated from either primary breast tumors collected at surgery (T-830, T-817, T-786, PDX-1915, PDX-1735 and PDX-1939) or from skin metastases (BM-156, BM-173) Prior treatments received by these TNBC patients are shown in **Supplementary table S1**. To generate PDXs, tumors were engrafted in the mammary fat pad of NSG (NOD-scid IL2Rgammanull) mice (Jackson Labs). Once tumors reached 2000mm^3^, tumors were collected and serially passaged in new NSG mice or preserved as live tissue in FBS 10% DMSO at −80°C for future engraftment.

PDXCs were generated from PDXs at passage P0-P1, in accordance with the Shlegel protocol^26^. Additional information can be found in the Supplementary methods.

Cells were routinely tested for Mycoplasma (MycoAlert PLUS Mycoplasma Detection Kit, Lonza, cat# LT07-518).

### Cell viability assay

Cells were seeded in 96-well plates and incubated at 37°C in 5% CO^2^ humidified atmosphere. Twenty-four hours later 100μL of F-medium containing the indicated supplements (**Supplementary Table S2**) was added. The media was removed 72h later and replaced with DMEM 10% Alamar Blue solution (Invitrogen cat# DAL1100). Fluorescence was read on a FLUOstar Optima (BMG Labtech), using 560-nm (Excitation) and 590-nm (Emission) filter settings.

For BM-156, given their poor adhesion, sulforhodamine B (SRB) assay was used instead of Alamar blue because with this method, cells were fixed with 10% trichloroacetic acid at 4°C before the removal of the medium at the end of the treatment (7 days). Cells were then rinsed with water before the addition of the SRB solution (0.2% SRB powder (Sigma cat#S1402) in 1% acetic acid) and incubated 30min at RT. The plates were rinsed with 1% acetic acid solution and Tris-Base pH 10.5 was added for 30min at RT. Absorbance was measured on a FLUOstar Optima.

Each condition was performed in triplicate. The percentage of cell viability was calculated by using the mean absorbance of the triplicate of a given condition normalized on the control or non-treated condition, which was set at 100% viability. Log dose response curves were plotted with GraphPad (version 5.0) and the IC50 for each condition was calculated by using the linear regression function, choosing a dose-response inhibition model (**Supplementary Table S2**).

### Combination index score

To determine if the combination of two drugs is synergistic, the Chou Talalay combination index (CI) score was used ^62^. The formulation can be found in Supplementary methods.

### Clonogenic assay

Cells were seeded in 6-well plates and treated with the indicated treatments (**Supplementary Table S2**) 24h later. After 14 days, cells were fixed with 10% formalin (Fisher cat#SF100-4) and stained with 0.5% crystal violet (Sigma, cat# HT90132-1L). Visible colonies were counted using a Gel Count colony counter (Oxford Optronix).

### PDX clinical trials

1-2mm^3^ pieces of live tissue were implanted into the mammary fat pad of 6–8-week NSG female mice for T-786, T-817 and BM-156 PDX models. Approximately 40 mice were engrafted, and tumor volume was measured every two days with an electronic caliper. Tumor volume was calculated by the formula (length × width^2^)/2. When tumors reached approximately 200mm^3^ mice were randomized: vehicle (0.9% saline water), carboplatin 20mg/kg intraperitoneal once a week, BAY1895344 40mg/kg once a week (gavage), carboplatin + BAY1895344, AZD6738 25mg/kg 5 days on 2 days off (gavage) and carboplatin + AZD6738. Tumor volume was measured twice weekly. Mice were evaluated for signs of toxicity by monitoring body weight and general body condition. Mice were euthanized with CO_2_ and isoflurane when tumors reached the maximal tumour size permitted (2000mm^3^) or body weight loss was >20% of the initial boy weight or there were other signs of animal distress or toxicity. The Studylog software (Studylog Systems) was used to facilitate data collection.

Smaller *in vivo* studies were also performed in additional PDX models (BM-173, PDX-1735, PDX-1886, PDX-1905, PDX-1915, PDX-1924, PDX-1939, PDX-1945, PDX-1971, PDX-1986, PDX-2076, PDX-2089). Mice were randomized into 3 treatment groups (∼3/group) only: vehicle, carboplatin 20mg/kg once weekly, and carboplatin + BAY1895344 40mg/kg once weekly. In cases where complete tumor response was observed, treatment was interrupted to allow tumor regrowth. Tumor growth inhibition formula can be found in Supplementary methods.

### Annexin V/PI apoptosis assay

Cells were seeded in a 6 well plate and treated 24h later as indicated **Supplementary Table S2**). After 72 hrs of treatment, floating dead cells were collected and combined with the trypsinized adherent cells. Apoptosis was measured by using the BD Pharmingen FITC Annexin V Apoptosis detection kit (cat. no. 556547) according to manufacturer’s instructions. Data were acquired on a FACS Canto II and analyzed with FACS Diva (BD Biosciences) or FlowJo software (Tree Star, Ashland, OR, USA).

### Genetic screens

To identify novel genes whose inhibition confers sensitivity to carboplatin, we performed RNAi-based genetic screens. We used an shRNA “kinome” library targeting 535 human kinases and kinase-related genes ^27,28^ and another shRNA library against ∼1,200 known target genes of clinically approved drugs. In brief, carboplatin resistant cells were transduced with the shRNA libraries at low MOI (Multiplicity Of Infection) (∼ 0.3) with a target of 1,000x coverage. Transduced cells were then selected in puromycin for 2 days and a time 0 (T0) was collected. Transduced cells were then treated with vehicle or an IC25 of carboplatin (23.5μM) for 14 days after which genomic DNA was extracted (time 1 or T1), and inserts were recovered with PCR amplification as described ^63^. The relative abundance of shRNAs in T0 and T1 were determined by next-generation sequencing and the relative abundance of shRNAs at T0 and T1 were analyzed by MAGeCK statistical software package (version 0.5.8)^64^ to identify the top candidates resensitizing cells to carboplatin^64^.

### siRNA and shRNA

Lentiviral transduction with low MOI was performed using the protocol described at http://carboplatin.broadinstitute.org/rnai/public/resources/protocols. Briefly, 2.5 × 10^6^ HEK293 T cells were seeded and transfected with indicated lentiviral constructs, the packaging (psPAX2) and envelope (pMD2.G) plasmid by CaCl2. Virus containing medium were collected for transduction of stable cell lines.

Following transduction, cells were selected with puromycin and/or blasticidin for 2–4 days and plated for downstream assays immediately after selection. Individual shRNA and ORF vectors were from the MISSION TRC library (Sigma), and ORF collections were developed by members of the ORFeome Collaboration (Sigma–TransOMIC), provided by the McGill Platform for Cellular Perturbation of the Goodman Cancer Research Centre. Sequences can be found in **Supplementary Table S**3.

For siRNA transfection, cells were seeded at 200 000 cells per well in a 6 well-plate, and 48h later, they were transfected with siRNA using lipofectamine RNAi/MAX (ThermoScientific cat# 13778075) following the manufacturer’s instructions. Information about the siRNAs can be found in **Supplementary Table S2**.

### Protein extraction and immunoblot analysis

Cells were lysed with RIPA buffer (ThermoScientific cat# 89900) supplemented with protease inhibitors aprotinin (10 μg/ml), leupeptin (10 μg/ml), phenylmethylsulfonyl fluoride (1 mM), and the phosphatase inhibitors Na_3_VO_4_ (1 mM) and NaF (5 mM) and incubated 15min on ice. Cells were centrifuged at 13 000 rpm at 4°C, the supernatant collected, and proteins quantified with Pierce BCA Protein Assay Kit (ThermoScientific cat# 23225). Membranes were blocked with 5% bovine serum albumin (BSA, Sigma-Aldrich cat# A1933), and incubated with primary antibodies (**Supplementary Table S4**) overnight at 4°C. Secondary antibodies were added the next day for 1h at RT. ECL chemiluminescence reagent to detect proteins (Sigma-Aldrich cat# GERPN2134 and cat# GERPN2235).

### Immunofluorescence

Cells were seeded on circular cover slips (Fisher Scientific, cat# 12545102P) and treated 24h later with drugs (**Supplementary Table S2**). Cells were then washed with PBS, fixed and permeabilized with PBS Triton X100 0.1% for 15min, washed with PBS and blocked with PBS 4% goat serum (Sigma-Aldrich cat# G9023). The cover slips were incubated for 1h30 with a mouse primary antibody against γH2AX (Millipore cat# JBW301), diluted in PBS 2% goat serum at 1:750, washed with PBS and then incubated in PBS 2% goat serum for 1h containing goat secondary antibody raised against mouse conjugated to Alexa-Fluor 594 (Invitrogen cat# A11020), DAPI (Invitrogen cat# D3571) and Alexa-Fluor 488 phalloidin (Invitrogen cat# A12379). The cells were washed before mounting the cover slips on slides. The slides were visualized using a Zeiss LSM800 confocal microscope with Airyscan. About 10 images per treatment group were unbiasedly acquired by selecting regions using the DAPI channel. Using Fiji software, a selection was made by contouring the cells using the phalloidin channel to then import this mask onto the Alexa-Fluor 594 channel to measure the mean fluorescence intensity. Each treatment groups contains at least 100 cells for each experiment, repeated 3 times.

### RT-qPCR

RNA extraction was done by using the RNeasy Mini Kit (QIAGEN cat#74104) with RNase A (QIAGEN Cat # 19101) and DNase (Qiagen Cat # 79254) according to the manufacturer’s instructions. Iscript cDNA synthesis kit (Biorad cat# 170-8891) was used to generate cDNAs according to the manufacturer’s instructions. Real time PCR was performed by using the SSOADV universal SYBR Green Supermix (BioRad, cat# 172-5272). ATR mRNA quantification was normalized by measuring GAPDH mRNA. Primers can be found in Supplementary Table S2.

### Cell cycle analysis

Cells were seeded in a 100mm culture dish. After 48h, cells were treated with the indicated drugs and incubated at 37°C for 24h. Cells were then washed, trypsinized, centrifuged, resuspended in PBS – 5mM EDTA and fixed for 30min with 100% ethanol while vortexing at low speed. Cells were pelleted and washed again with PBS – 5mM EDTA and resuspended at 1×10^6^ cells/mL in PBS with 50ug/mL propidium iodide and 20μg/ml RNAseA DNAse free. Cells were finally incubated at 37°C in the dark for 30min and analyzed on a FACS Canto II instrument (Becton Dickinson). Results were analyzed using ModFit LT (Verity Software House, version 4.1.7).

### RNA-sequencing of PDXCs

RNA was extracted using Qiagen RNeasy Mini kit (QIAGEN cat. #74104) according to the manufacturer’s instructions. Triplicates for each condition were sequenced using the Illumina NextSeq500 platform 1 × 75 cycle High Output. RNA sequencing libraries were prepared from total RNA using poly(A) enrichment. Data analysis information can be found in Supplementary methods.

### Statistics

Statistical analyses were performed using GraphPad software 5.0 (Prism). Data in figures are represented as mean ± SEM. For the bar graphs and the dot plot, statistical differences were assessed using Student’s *t* test to compare 2 groups of interest. For tumor growth, statistical significance was assessed using Student’s *t* test to compare 2 groups of interest at the end of the study. Mantel-Cox analysis of Kaplan-Meier curves was performed to analyze statistical differences in survival. Significance was considered with P value under 0.05: *P < 0.05, ***P < 0.01, ***P < 0.001.

### Study approval

The study was conducted in accordance with the Declaration of Helsinki and approved by the Institutional Review Board (or Ethics Committee) of the Jewish General Hospital. Patients consented to sample collection as part of the breast biobank of the Jewish General Hospital (protocol #05-006). PDX and PDXCs generation was performed as part of protocol #14-168 approved by the JGH review ethics board. All animal studies were conducted in accordance with the regulations formulated by McGill University’s Animal Care and Use Committee.

## Data availability

Data were generated by the authors and available on request.

## Models availability

To promote transparency and ensure reproducibility, we are committed to sharing our PDX and PDXC models. This is subject to the approval of the institutional review ethics board.

## Acknowledgement.

This research was supported by the Cancer Research Society, Canadian Institutes of Health Research and an Oncopole EMC2 grant to MB. AA-M is supported by the Guerrera Familly Cancer Scientist Award. We thank Le Réseau Recherche Cancer, McPeak Sirois Consortium, the Quebec Breast Cancer Foundation and the Quebec Cancer Consortium for supporting the JGH Breast Cancer Biobank and the Breast Cancer Functional Genomics Group (BCFGG). We also thank Genome Quebec and the Institute of Research in Immunology and Cancer (IRIC) for sequencing services. We gratefully thank REPARE Therapeutics for supplying us with RP-6306.

## References

1. Cortazar, P., et al. Pathological complete response and long-term clinical benefit in breast cancer: the CTNeoBC pooled analysis. Lancet 384, 164–172 (2014).

2. Ignatov, A., Eggemann, H., Burger, E. & Ignatov, T. Patterns of breast cancer relapse in accordance to biological subtype. J Cancer Res Clin Oncol 144, 1347–1355 (2018).

3. Rakha, E.A., et al. Prognostic markers in triple-negative breast cancer. Cancer 109, 25–32 (2007).

4. Mandapati, A. & Lukong, K.E. Triple negative breast cancer: approved treatment options and their mechanisms of action. J Cancer Res Clin Oncol 149, 3701–3719 (2023).

5. Poggio, F., et al. Platinum-based neoadjuvant chemotherapy in triple-negative breast cancer: a systematic review and meta-analysis. Ann Oncol 29, 1497–1508 (2018).

6. Zhou, J., et al. The Drug-Resistance Mechanisms of Five Platinum-Based Antitumor Agents. Front Pharmacol 11, 343 (2020).

7. Moens, S., et al. The mitotic checkpoint is a targetable vulnerability of carboplatin-resistant triple negative breast cancers. Sci Rep 11, 3176 (2021).

8. Lord, C.J. & Ashworth, A. BRCAness revisited. Nat Rev Cancer 16, 110–120 (2016).

9. Saldivar, J.C., et al. An intrinsic S/G2 checkpoint enforced by ATR. Science 361, 806–810 (2018).

10. Xiao, Z., et al. Chk1 mediates S and G2 arrests through Cdc25A degradation in response to DNA-damaging agents. J Biol Chem 278, 21767–21773 (2003).

11. Bradbury, A., Hall, S., Curtin, N. & Drew, Y. Targeting ATR as Cancer Therapy: A new era for synthetic lethality and synergistic combinations? Pharmacol Ther 207, 107450 (2020).

12. Lecona, E. & Fernandez-Capetillo, O. Targeting ATR in cancer. Nat Rev Cancer 18, 586–595 (2018).

13. Barnieh, F.M., Loadman, P.M. & Falconer, R.A. Progress towards a clinically-successful ATR inhibitor for cancer therapy. Curr Res Pharmacol Drug Discov 2, 100017 (2021).

14. Ngoi, N.Y.L., et al. Targeting ATR in patients with cancer. Nat Rev Clin Oncol 21, 278–293 (2024).

15. Kwok, M., et al. ATR inhibition induces synthetic lethality and overcomes chemoresistance in TP53- or ATM-defective chronic lymphocytic leukemia cells. Blood 127, 582–595 (2016).

16. Shapiro, G.I., et al. Phase 1 study of the ATR inhibitor berzosertib in combination with cisplatin in patients with advanced solid tumours. Br J Cancer 125, 520–527 (2021).

17. Yap, T.A., et al. Ceralasertib (AZD6738), an Oral ATR Kinase Inhibitor, in Combination with Carboplatin in Patients with Advanced Solid Tumors: A Phase I Study. Clin Cancer Res 27, 5213–5224 (2021).

18. Heyza, J.R., et al. ATR inhibition overcomes platinum tolerance associated with ERCC1- and p53-deficiency by inducing replication catastrophe. NAR Cancer 5, zcac045 (2023).

19. Konig, P., et al. SLFN11 and ATR as targets for overcoming cisplatin resistance in ovarian cancer cells. Biochim Biophys Acta Mol Basis Dis 1870, 167448 (2024).

20. Fattaey, A. & Booher, R.N. Myt1: a Wee1-type kinase that phosphorylates Cdc2 on residue Thr14. Prog Cell Cycle Res 3, 233–240 (1997).

21. Chow, J.P., Poon, R.Y. & Ma, H.T. Inhibitory phosphorylation of cyclin-dependent kinase 1 as a compensatory mechanism for mitosis exit. Mol Cell Biol 31, 1478–1491 (2011).

22. Chow, J.P. & Poon, R.Y. The CDK1 inhibitory kinase MYT1 in DNA damage checkpoint recovery. Oncogene 32, 4778–4788 (2013).

23. Li, M., et al. Low Molecular Weight Cyclin E Confers a Vulnerability to PKMYT1 Inhibition in Triple-Negative Breast Cancer. Cancer Res (2024).

24. Chen, A., et al. PKMYT1 is a Marker of Treatment Response and a Therapeutic Target for CDK4/6 Inhibitor-Resistance in ER+ Breast Cancer. Mol Cancer Ther (2024).

25. Lang, F., et al. Abrogation of the G2/M checkpoint as a chemosensitization approach for alkylating agents. Neuro Oncol 26, 1083–1096 (2024).

26. Liu, X., et al. Conditional reprogramming and long-term expansion of normal and tumor cells from human biospecimens. Nat Protoc 12, 439–451 (2017).

27. Kong, T., et al. CD44 Promotes PD-L1 Expression and Its Tumor-Intrinsic Function in Breast and Lung Cancers. Cancer Res 80, 444–457 (2020).

28. Prahallad, A., et al. Unresponsiveness of colon cancer to BRAF(V600E) inhibition through feedback activation of EGFR. Nature 483, 100–103 (2012).

29. Liu, Q., et al. Chk1 is an essential kinase that is regulated by Atr and required for the G(2)/M DNA damage checkpoint. Genes Dev 14, 1448–1459 (2000).

30. Zhao, H. & Piwnica-Worms, H. ATR-mediated checkpoint pathways regulate phosphorylation and activation of human Chk1. Mol Cell Biol 21, 4129–4139 (2001).

31. Smits, V.A., Reaper, P.M. & Jackson, S.P. Rapid PIKK-dependent release of Chk1 from chromatin promotes the DNA-damage checkpoint response. Curr Biol 16, 150–159 (2006).

32. Toledo, L.I., et al. ATR prohibits replication catastrophe by preventing global exhaustion of RPA. Cell 155, 1088–1103 (2013).

33. Vassin, V.M., Anantha, R.W., Sokolova, E., Kanner, S. & Borowiec, J.A. Human RPA phosphorylation by ATR stimulates DNA synthesis and prevents ssDNA accumulation during DNA-replication stress. J Cell Sci 122, 4070–4080 (2009).

34. Ashley, A.K., et al. DNA-PK phosphorylation of RPA32 Ser4/Ser8 regulates replication stress checkpoint activation, fork restart, homologous recombination and mitotic catastrophe. DNA Repair (Amst) 21, 131–139 (2014).

35. Mah, L.J., El-Osta, A. & Karagiannis, T.C. gammaH2AX: a sensitive molecular marker of DNA damage and repair. Leukemia 24, 679–686 (2010).

36. Ziv, Y., et al. Chromatin relaxation in response to DNA double-strand breaks is modulated by a novel ATM- and KAP-1 dependent pathway. Nat Cell Biol 8, 870–876 (2006).

37. Moeglin, E., et al. Uniform Widespread Nuclear Phosphorylation of Histone H2AX Is an Indicator of Lethal DNA Replication Stress. Cancers (Basel) 11(2019).

38. Sorensen, C.S., et al. Chk1 regulates the S phase checkpoint by coupling the physiological turnover and ionizing radiation-induced accelerated proteolysis of Cdc25A. Cancer Cell 3, 247–258 (2003).

39. Sanchez, Y., et al. Conservation of the Chk1 checkpoint pathway in mammals: linkage of DNA damage to Cdk regulation through Cdc25. Science 277, 1497–1501 (1997).

40. Castedo, M., et al. Cell death by mitotic catastrophe: a molecular definition. Oncogene 23, 2825–2837 (2004).

41. Lachmann, A., et al. ChEA: transcription factor regulation inferred from integrating genome-wide ChIP-X experiments. Bioinformatics 26, 2438–2444 (2010).

42. Lindqvist, A., Rodriguez-Bravo, V. & Medema, R.H. The decision to enter mitosis: feedback and redundancy in the mitotic entry network. J Cell Biol 185, 193–202 (2009).

43. Gavet, O. & Pines, J. Progressive activation of CyclinB1-Cdk1 coordinates entry to mitosis. Dev Cell 18, 533–543 (2010).

44. Liu, F., Stanton, J.J., Wu, Z. & Piwnica-Worms, H. The human Myt1 kinase preferentially phosphorylates Cdc2 on threonine 14 and localizes to the endoplasmic reticulum and Golgi complex. Mol Cell Biol 17, 571–583 (1997).

45. Mueller, P.R., Coleman, T.R., Kumagai, A. & Dunphy, W.G. Myt1: a membrane-associated inhibitory kinase that phosphorylates Cdc2 on both threonine-14 and tyrosine-15. Science 270, 86–90 (1995).

46. Elbaek, C.R., Petrosius, V. & Sorensen, C.S. WEE1 kinase limits CDK activities to safeguard DNA replication and mitotic entry. Mutat Res 819-820, 111694 (2020).

47. Booher, R.N., Holman, P.S. & Fattaey, A. Human Myt1 is a cell cycle-regulated kinase that inhibits Cdc2 but not Cdk2 activity. J Biol Chem 272, 22300–22306 (1997).

48. Nakajima, H., Toyoshima-Morimoto, F., Taniguchi, E. & Nishida, E. Identification of a consensus motif for Plk (Polo-like kinase) phosphorylation reveals Myt1 as a Plk1 substrate. J Biol Chem 278, 25277–25280 (2003).

49. Okumura, E., et al. Akt inhibits Myt1 in the signalling pathway that leads to meiotic G2/M-phase transition. Nat Cell Biol 4, 111–116 (2002).

50. Villeneuve, J., Scarpa, M., Ortega-Bellido, M. & Malhotra, V. MEK1 inactivates Myt1 to regulate Golgi membrane fragmentation and mitotic entry in mammalian cells. EMBO J 32, 72–85 (2013).

51. Ruiz, E.J., Vilar, M. & Nebreda, A.R. A two-step inactivation mechanism of Myt1 ensures CDK1/cyclin B activation and meiosis I entry. Curr Biol 20, 717–723 (2010).

52. Szychowski, J., et al. Discovery of an Orally Bioavailable and Selective PKMYT1 Inhibitor, RP-6306. J Med Chem 65, 10251–10284 (2022).

53. Gallo, D., et al. CCNE1 amplification is synthetic lethal with PKMYT1 kinase inhibition. Nature 604, 749–756 (2022).

54. Yap, T.A., et al. Camonsertib in DNA damage response-deficient advanced solid tumors: phase 1 trial results. Nat Med 29, 1400–1411 (2023).

55. Hustedt, N., et al. A consensus set of genetic vulnerabilities to ATR inhibition. Open Biol 9, 190156 (2019).

56. Williamson, C.T., et al. ATR inhibitors as a synthetic lethal therapy for tumours deficient in ARID1A. Nat Commun 7, 13837 (2016).

57. Harold, J., et al. Elimusertib (BAY1895344), a novel ATR inhibitor, demonstrates in vivo activity in ATRX mutated models of uterine leiomyosarcoma. Gynecol Oncol 168, 157–165 (2023).

58. Savage, P., et al. Chemogenomic profiling of breast cancer patient-derived xenografts reveals targetable vulnerabilities for difficult-to-treat tumors. Commun Biol 3, 310 (2020).

59. Martin, L.P., Hamilton, T.C. & Schilder, R.J. Platinum resistance: the role of DNA repair pathways. Clin Cancer Res 14, 1291–1295 (2008).

60. Forgie, B.N., Prakash, R. & Telleria, C.M. Revisiting the Anti-Cancer Toxicity of Clinically Approved Platinating Derivatives. Int J Mol Sci 23(2022).

61. Peng, C.Y., et al. Mitotic and G2 checkpoint control: regulation of 14-3-3 protein binding by phosphorylation of Cdc25C on serine-216. Science 277, 1501–1505 (1997).

62. Chou, T.C. & Talalay, P. Quantitative analysis of dose-effect relationships: the combined effects of multiple drugs or enzyme inhibitors. Adv Enzyme Regul 22, 27–55 (1984).

63. Sun, C., et al. Reversible and adaptive resistance to BRAF(V600E) inhibition in melanoma. Nature 508, 118–122 (2014).

64. Li, W., et al. MAGeCK enables robust identification of essential genes from genome-scale CRISPR/Cas9 knockout screens. Genome Biol 15, 554 (2014).

