## Supplementary figures and tables for "ATR and PKMYT1 inhibition re-sensitize a subset of TNBC patient-derived models to carboplatin inducing mitotic catastrophe"

### Slide 1
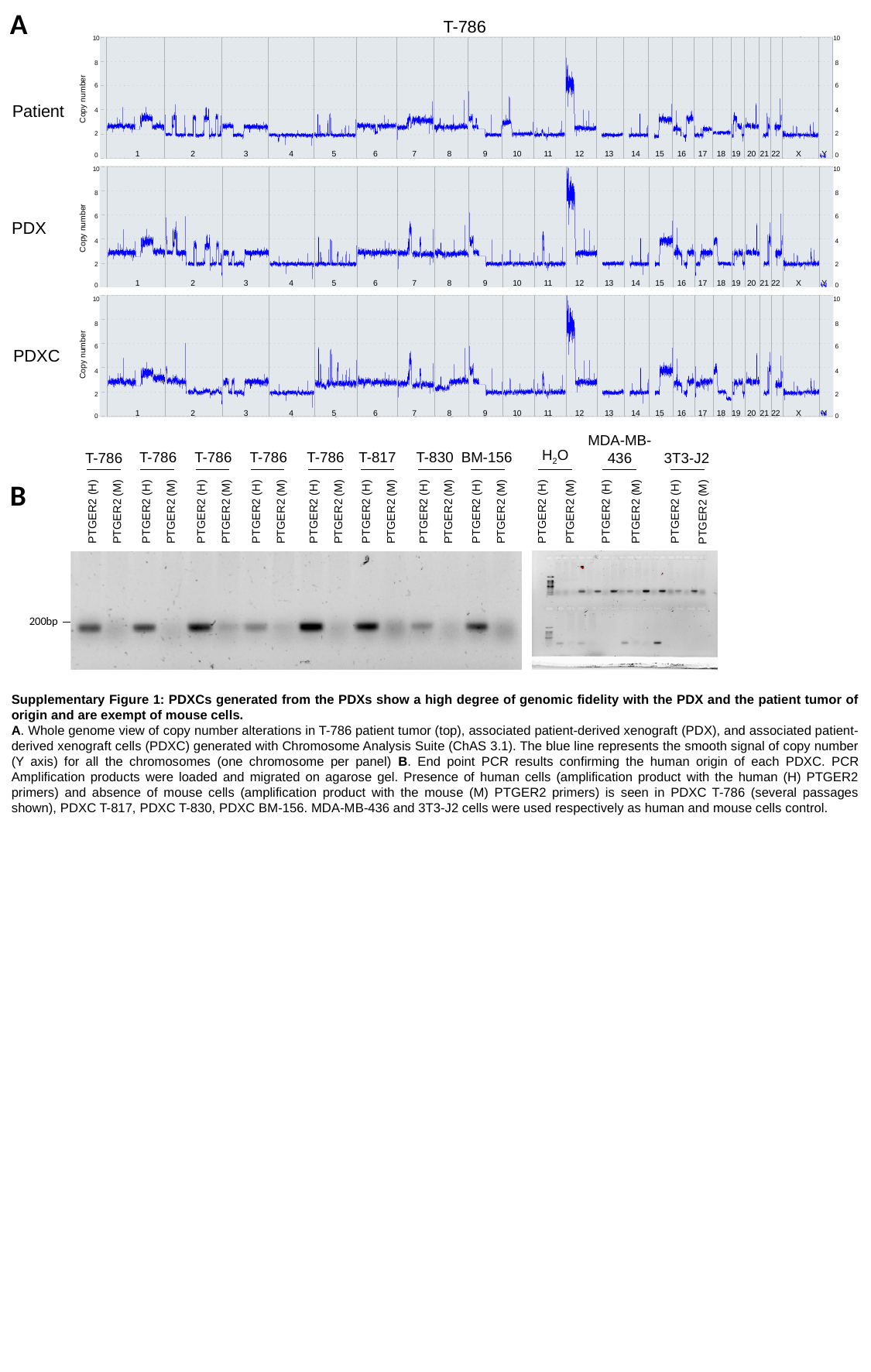

A
T-786
Patient
PDX
PDXC
10
8
6
4
2
0
10
8
6
4
2
0
Copy number
1
2
3
4
5
6
7
8
9
10
11
12
13
14
15
16
17
18
19
20
21
22
X
Y
10
8
6
4
2
0
10
8
6
4
2
0
Copy number
1
2
3
4
5
6
7
8
9
10
11
12
13
14
15
16
17
18
19
20
21
22
X
Y
10
8
6
4
2
0
10
8
6
4
2
0
Copy number
1
2
3
4
5
6
7
8
9
10
11
12
13
14
15
16
17
18
19
20
21
22
X
Y
MDA-MB-
436
H_2_O
T-817
T-830
BM-156
T-786
T-786
T-786
T-786
3T3-J2
T-786
PTGER2 (H)
PTGER2 (M)
PTGER2 (H)
PTGER2 (M)
PTGER2 (H)
PTGER2 (M)
PTGER2 (H)
PTGER2 (M)
PTGER2 (H)
PTGER2 (M)
PTGER2 (H)
PTGER2 (M)
PTGER2 (H)
PTGER2 (M)
PTGER2 (H)
PTGER2 (M)
PTGER2 (H)
PTGER2 (M)
PTGER2 (H)
PTGER2 (M)
PTGER2 (H)
PTGER2 (M)
200bp
B
Supplementary Figure 1: PDXCs generated from the PDXs show a high degree of genomic fidelity with the PDX and the patient tumor of origin and are exempt of mouse cells.
A. Whole genome view of copy number alterations in T-786 patient tumor (top), associated patient-derived xenograft (PDX), and associated patient-derived xenograft cells (PDXC) generated with Chromosome Analysis Suite (ChAS 3.1). The blue line represents the smooth signal of copy number (Y axis) for all the chromosomes (one chromosome per panel) B. End point PCR results confirming the human origin of each PDXC. PCR Amplification products were loaded and migrated on agarose gel. Presence of human cells (amplification product with the human (H) PTGER2 primers) and absence of mouse cells (amplification product with the mouse (M) PTGER2 primers) is seen in PDXC T-786 (several passages shown), PDXC T-817, PDXC T-830, PDXC BM-156. MDA-MB-436 and 3T3-J2 cells were used respectively as human and mouse cells control.

### Slide 2
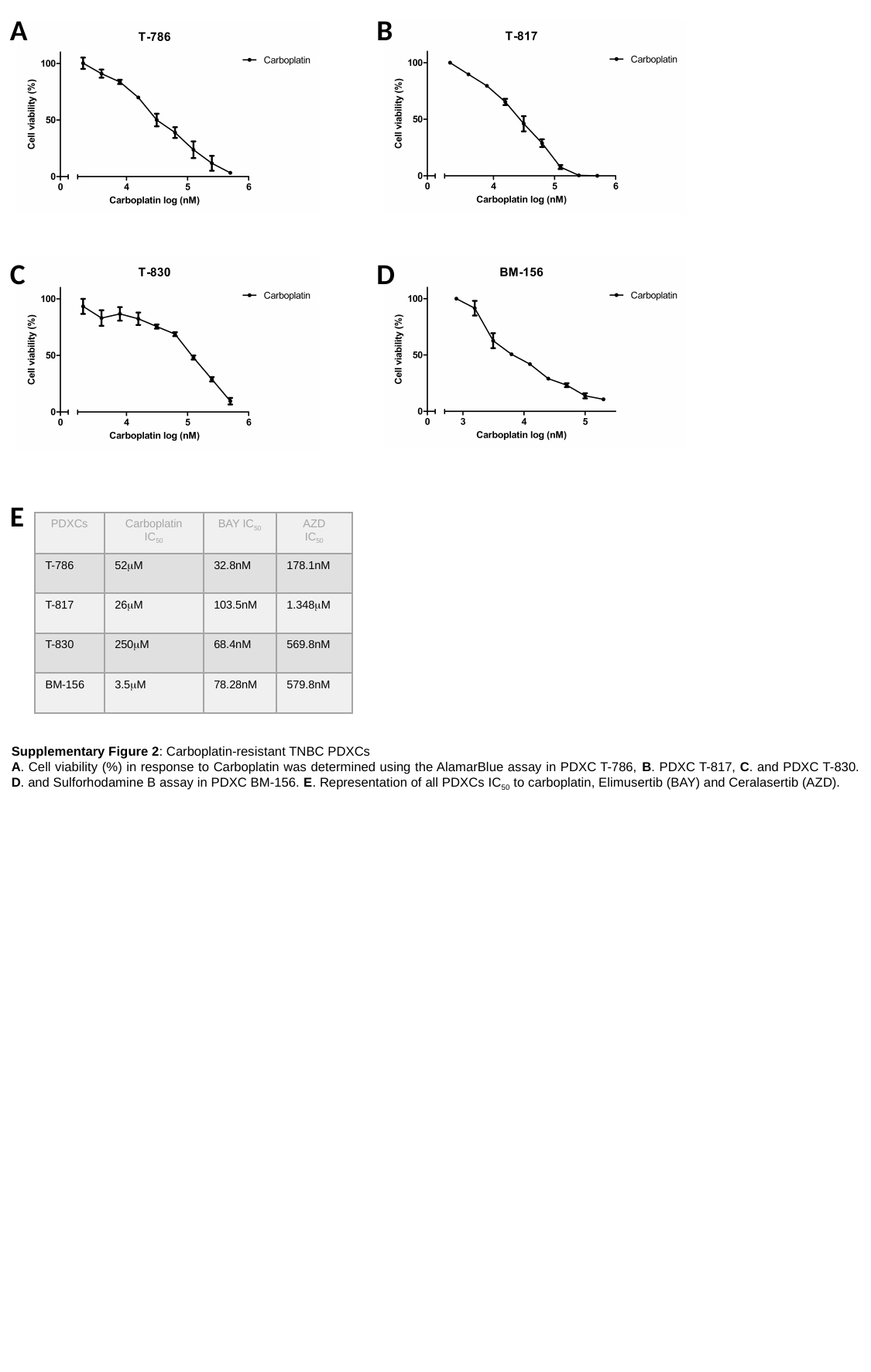

A
B
C
D
E
| PDXCs | Carboplatin IC_50_ | BAY IC_50_ | AZD IC_50_ |
| --- | --- | --- | --- |
| T-786 | 52μM | 32.8nM | 178.1nM |
| T-817 | 26μM | 103.5nM | 1.348μM |
| T-830 | 250μM | 68.4nM | 569.8nM |
| BM-156 | 3.5μM | 78.28nM | 579.8nM |
Supplementary Figure 2: Carboplatin-resistant TNBC PDXCs
A. Cell viability (%) in response to Carboplatin was determined using the AlamarBlue assay in PDXC T-786, B. PDXC T-817, C. and PDXC T-830. D. and Sulforhodamine B assay in PDXC BM-156. E. Representation of all PDXCs IC_50_ to carboplatin, Elimusertib (BAY) and Ceralasertib (AZD).

### Slide 3
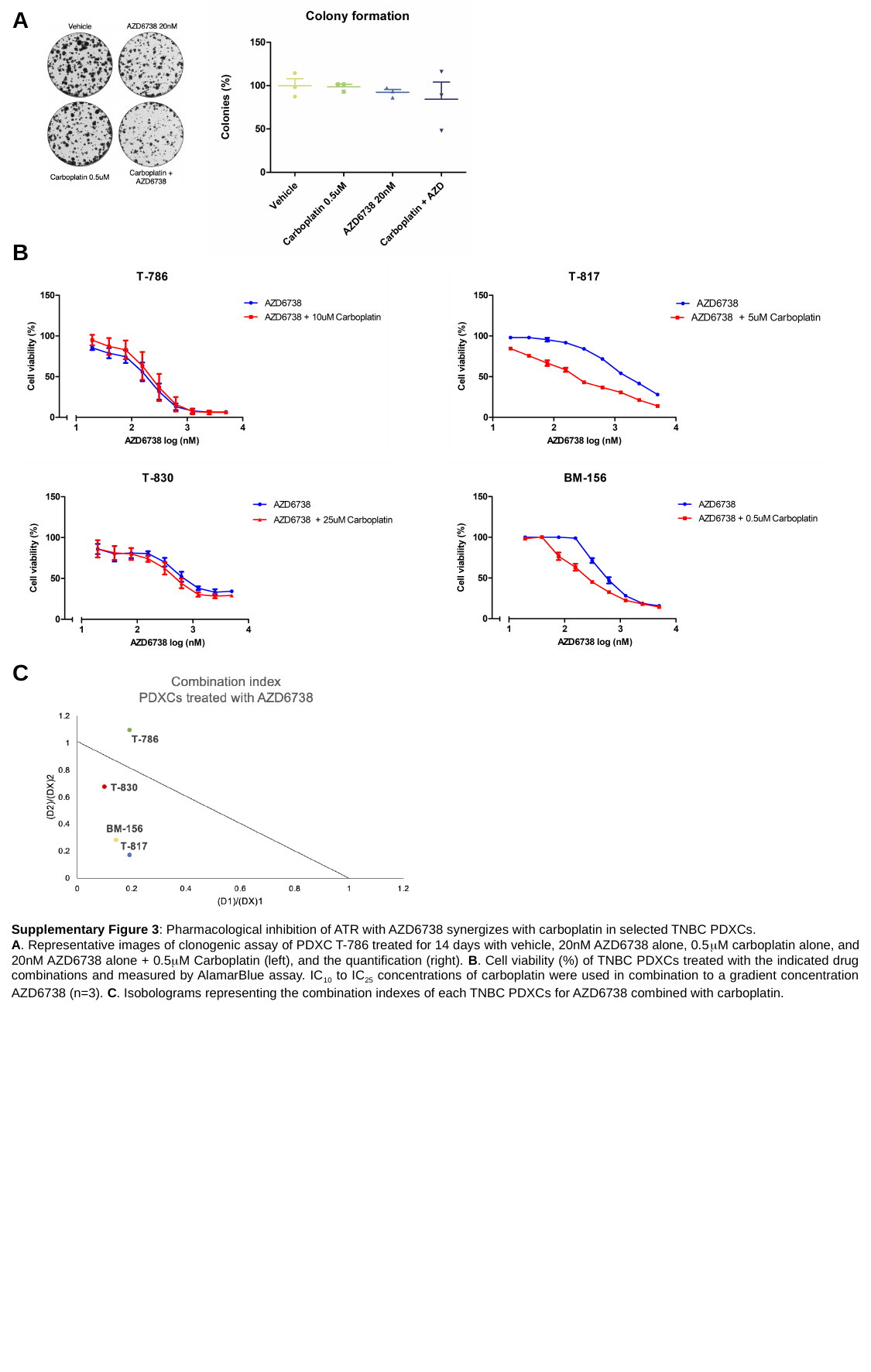

A
B
C
Supplementary Figure 3: Pharmacological inhibition of ATR with AZD6738 synergizes with carboplatin in selected TNBC PDXCs.
A. Representative images of clonogenic assay of PDXC T-786 treated for 14 days with vehicle, 20nM AZD6738 alone, 0.5μM carboplatin alone, and 20nM AZD6738 alone + 0.5μM Carboplatin (left), and the quantification (right). B. Cell viability (%) of TNBC PDXCs treated with the indicated drug combinations and measured by AlamarBlue assay. IC_10_ to IC_25_ concentrations of carboplatin were used in combination to a gradient concentration AZD6738 (n=3). C. Isobolograms representing the combination indexes of each TNBC PDXCs for AZD6738 combined with carboplatin.

### Slide 4
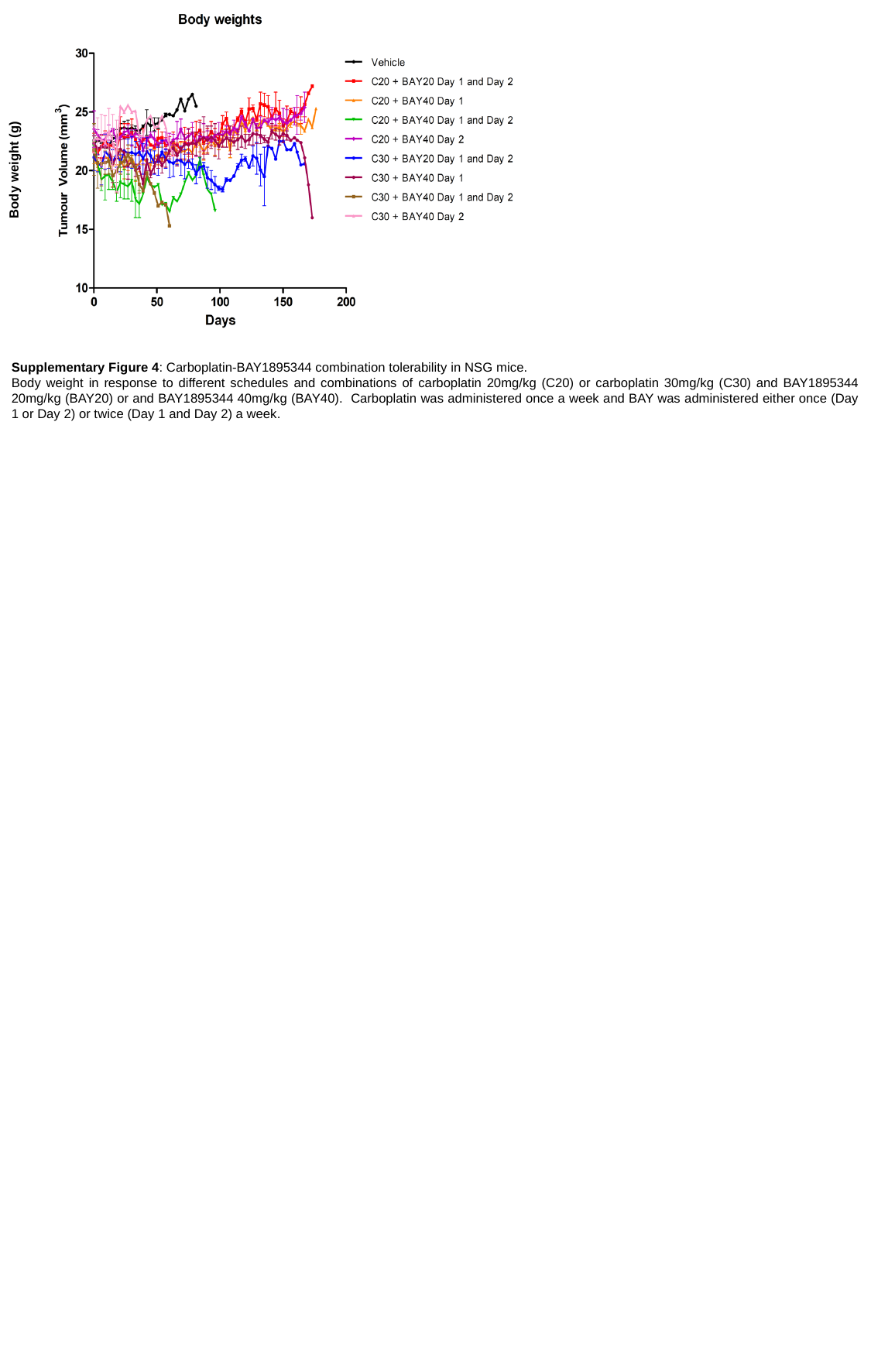

Body weight (g)
Supplementary Figure 4: Carboplatin-BAY1895344 combination tolerability in NSG mice.
Body weight in response to different schedules and combinations of carboplatin 20mg/kg (C20) or carboplatin 30mg/kg (C30) and BAY1895344 20mg/kg (BAY20) or and BAY1895344 40mg/kg (BAY40). Carboplatin was administered once a week and BAY was administered either once (Day 1 or Day 2) or twice (Day 1 and Day 2) a week.

### Slide 5
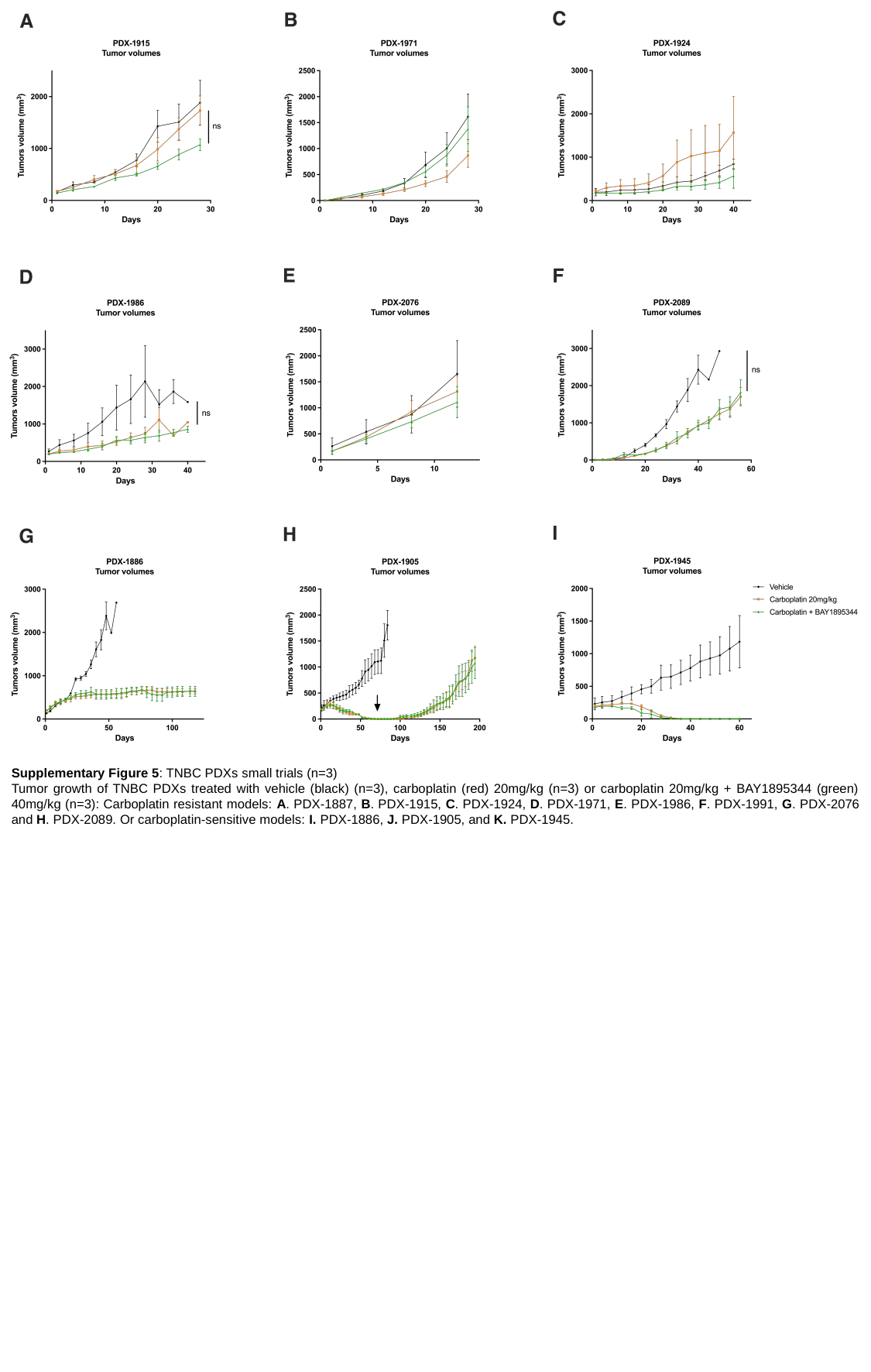

Supplementary Figure 5: TNBC PDXs small trials (n=3)
Tumor growth of TNBC PDXs treated with vehicle (black) (n=3), carboplatin (red) 20mg/kg (n=3) or carboplatin 20mg/kg + BAY1895344 (green) 40mg/kg (n=3): Carboplatin resistant models: A. PDX-1887, B. PDX-1915, C. PDX-1924, D. PDX-1971, E. PDX-1986, F. PDX-1991, G. PDX-2076 and H. PDX-2089. Or carboplatin-sensitive models: I. PDX-1886, J. PDX-1905, and K. PDX-1945.

### Slide 6
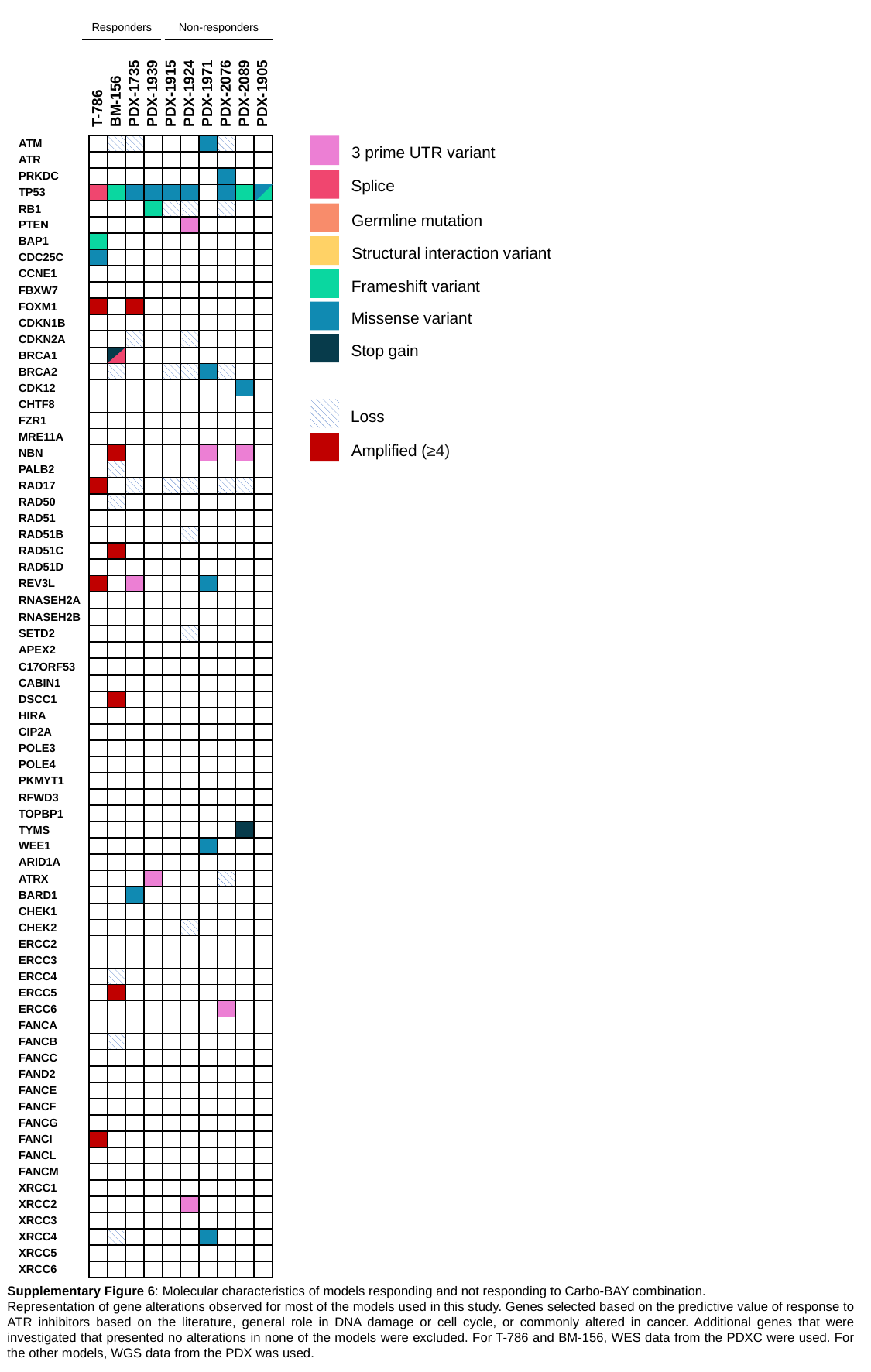

Responders
Non-responders
| | T-786 | BM-156 | PDX-1735 | PDX-1939 | PDX-1915 | PDX-1924 | PDX-1971 | PDX-2076 | PDX-2089 | PDX-1905 |
| --- | --- | --- | --- | --- | --- | --- | --- | --- | --- | --- |
| ATM | | | | | | | | | | |
| ATR | | | | | | | | | | |
| PRKDC | | | | | | | | | | |
| TP53 | | | | | | | | | | |
| RB1 | | | | | | | | | | |
| PTEN | | | | | | | | | | |
| BAP1 | | | | | | | | | | |
| CDC25C | | | | | | | | | | |
| CCNE1 | | | | | | | | | | |
| FBXW7 | | | | | | | | | | |
| FOXM1 | | | | | | | | | | |
| CDKN1B | | | | | | | | | | |
| CDKN2A | | | | | | | | | | |
| BRCA1 | | | | | | | | | | |
| BRCA2 | | | | | | | | | | |
| CDK12 | | | | | | | | | | |
| CHTF8 | | | | | | | | | | |
| FZR1 | | | | | | | | | | |
| MRE11A | | | | | | | | | | |
| NBN | | | | | | | | | | |
| PALB2 | | | | | | | | | | |
| RAD17 | | | | | | | | | | |
| RAD50 | | | | | | | | | | |
| RAD51 | | | | | | | | | | |
| RAD51B | | | | | | | | | | |
| RAD51C | | | | | | | | | | |
| RAD51D | | | | | | | | | | |
| REV3L | | | | | | | | | | |
| RNASEH2A | | | | | | | | | | |
| RNASEH2B | | | | | | | | | | |
| SETD2 | | | | | | | | | | |
| APEX2 | | | | | | | | | | |
| C17ORF53 | | | | | | | | | | |
| CABIN1 | | | | | | | | | | |
| DSCC1 | | | | | | | | | | |
| HIRA | | | | | | | | | | |
| CIP2A | | | | | | | | | | |
| POLE3 | | | | | | | | | | |
| POLE4 | | | | | | | | | | |
| PKMYT1 | | | | | | | | | | |
| RFWD3 | | | | | | | | | | |
| TOPBP1 | | | | | | | | | | |
| TYMS | | | | | | | | | | |
| WEE1 | | | | | | | | | | |
| ARID1A | | | | | | | | | | |
| ATRX | | | | | | | | | | |
| BARD1 | | | | | | | | | | |
| CHEK1 | | | | | | | | | | |
| CHEK2 | | | | | | | | | | |
| ERCC2 | | | | | | | | | | |
| ERCC3 | | | | | | | | | | |
| ERCC4 | | | | | | | | | | |
| ERCC5 | | | | | | | | | | |
| ERCC6 | | | | | | | | | | |
| FANCA | | | | | | | | | | |
| FANCB | | | | | | | | | | |
| FANCC | | | | | | | | | | |
| FAND2 | | | | | | | | | | |
| FANCE | | | | | | | | | | |
| FANCF | | | | | | | | | | |
| FANCG | | | | | | | | | | |
| FANCI | | | | | | | | | | |
| FANCL | | | | | | | | | | |
| FANCM | | | | | | | | | | |
| XRCC1 | | | | | | | | | | |
| XRCC2 | | | | | | | | | | |
| XRCC3 | | | | | | | | | | |
| XRCC4 | | | | | | | | | | |
| XRCC5 | | | | | | | | | | |
| XRCC6 | | | | | | | | | | |
3 prime UTR variant
Splice
Germline mutation
Structural interaction variant
Frameshift variant
Missense variant
Stop gain
Loss
Amplified (≥4)
Supplementary Figure 6: Molecular characteristics of models responding and not responding to Carbo-BAY combination.
Representation of gene alterations observed for most of the models used in this study. Genes selected based on the predictive value of response to ATR inhibitors based on the literature, general role in DNA damage or cell cycle, or commonly altered in cancer. Additional genes that were investigated that presented no alterations in none of the models were excluded. For T-786 and BM-156, WES data from the PDXC were used. For the other models, WGS data from the PDX was used.

### Slide 7
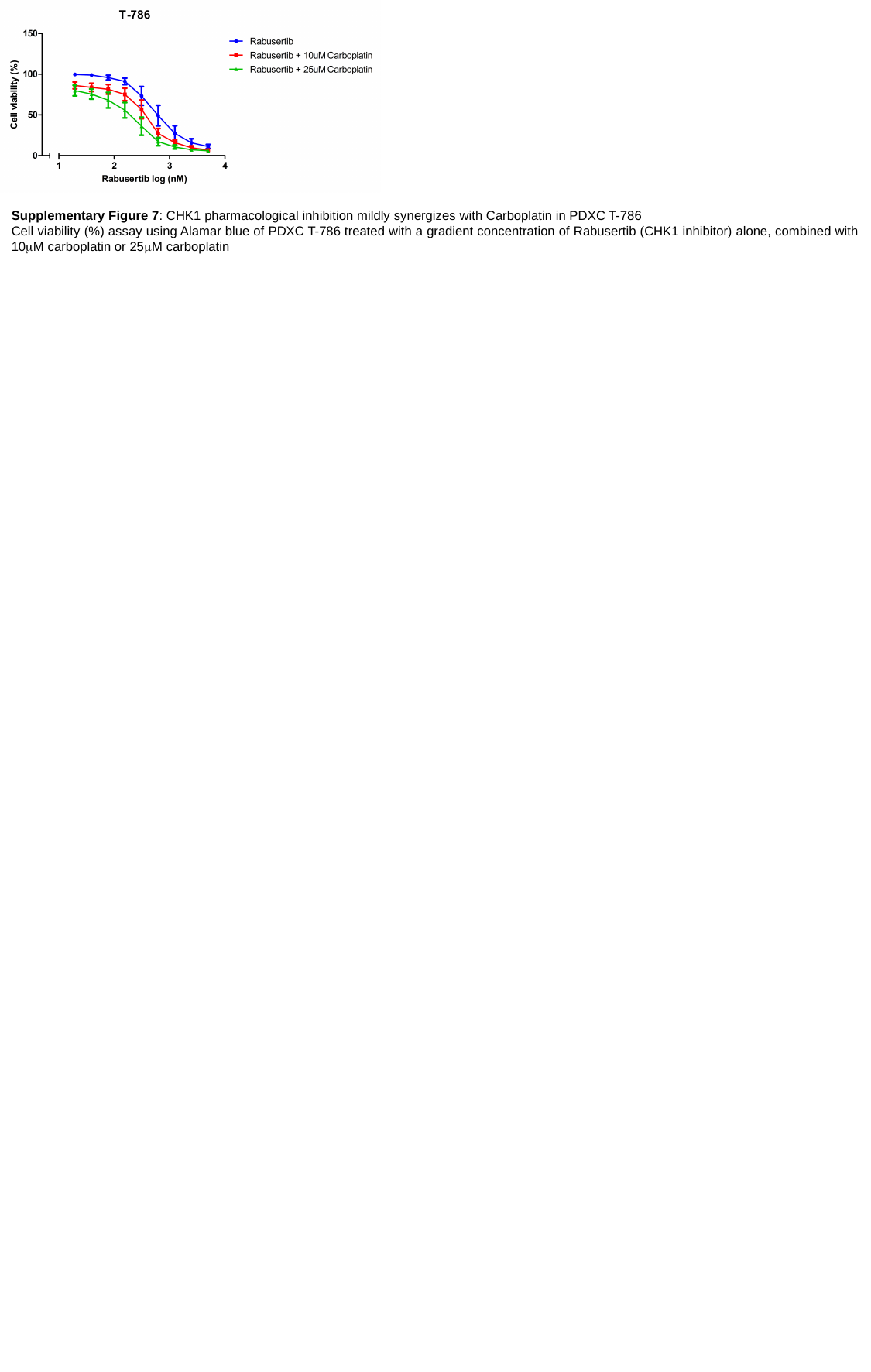

Supplementary Figure 7: CHK1 pharmacological inhibition mildly synergizes with Carboplatin in PDXC T-786
Cell viability (%) assay using Alamar blue of PDXC T-786 treated with a gradient concentration of Rabusertib (CHK1 inhibitor) alone, combined with 10μM carboplatin or 25μM carboplatin

### Slide 8
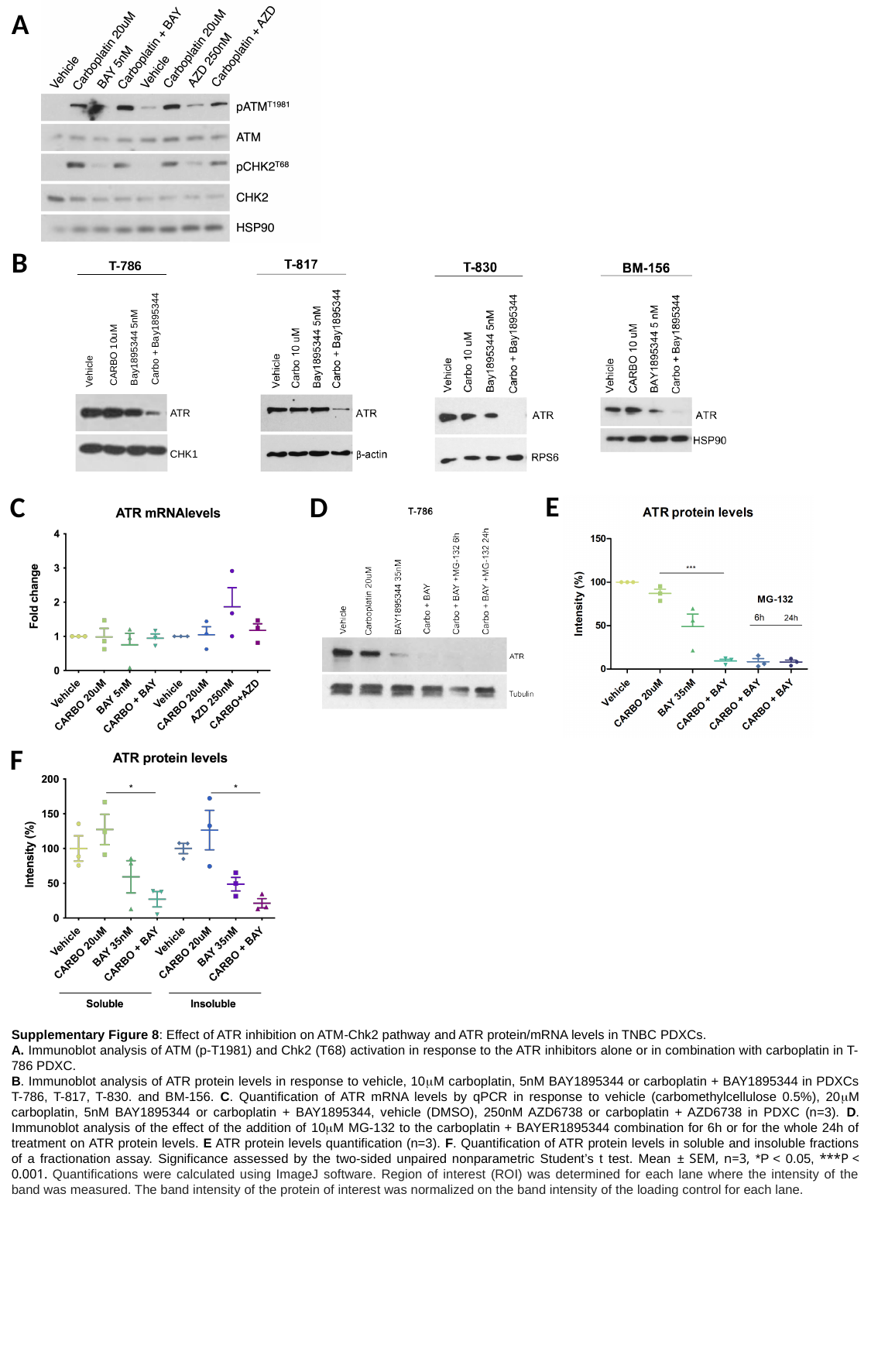

A
B
T-786
Carbo + Bay1895344
Bay1895344 5nM
CARBO 10uM
Vehicle
ATR
CHK1
E
C
D
F
Supplementary Figure 8: Effect of ATR inhibition on ATM-Chk2 pathway and ATR protein/mRNA levels in TNBC PDXCs.
A. Immunoblot analysis of ATM (p-T1981) and Chk2 (T68) activation in response to the ATR inhibitors alone or in combination with carboplatin in T-786 PDXC.
B. Immunoblot analysis of ATR protein levels in response to vehicle, 10μM carboplatin, 5nM BAY1895344 or carboplatin + BAY1895344 in PDXCs T-786, T-817, T-830. and BM-156. C. Quantification of ATR mRNA levels by qPCR in response to vehicle (carbomethylcellulose 0.5%), 20μM carboplatin, 5nM BAY1895344 or carboplatin + BAY1895344, vehicle (DMSO), 250nM AZD6738 or carboplatin + AZD6738 in PDXC (n=3). D. Immunoblot analysis of the effect of the addition of 10μM MG-132 to the carboplatin + BAYER1895344 combination for 6h or for the whole 24h of treatment on ATR protein levels. E ATR protein levels quantification (n=3). F. Quantification of ATR protein levels in soluble and insoluble fractions of a fractionation assay. Significance assessed by the two-sided unpaired nonparametric Student’s t test. Mean ± SEM, n=3, *P < 0.05, ***P < 0.001. Quantifications were calculated using ImageJ software. Region of interest (ROI) was determined for each lane where the intensity of the band was measured. The band intensity of the protein of interest was normalized on the band intensity of the loading control for each lane.

### Slide 9
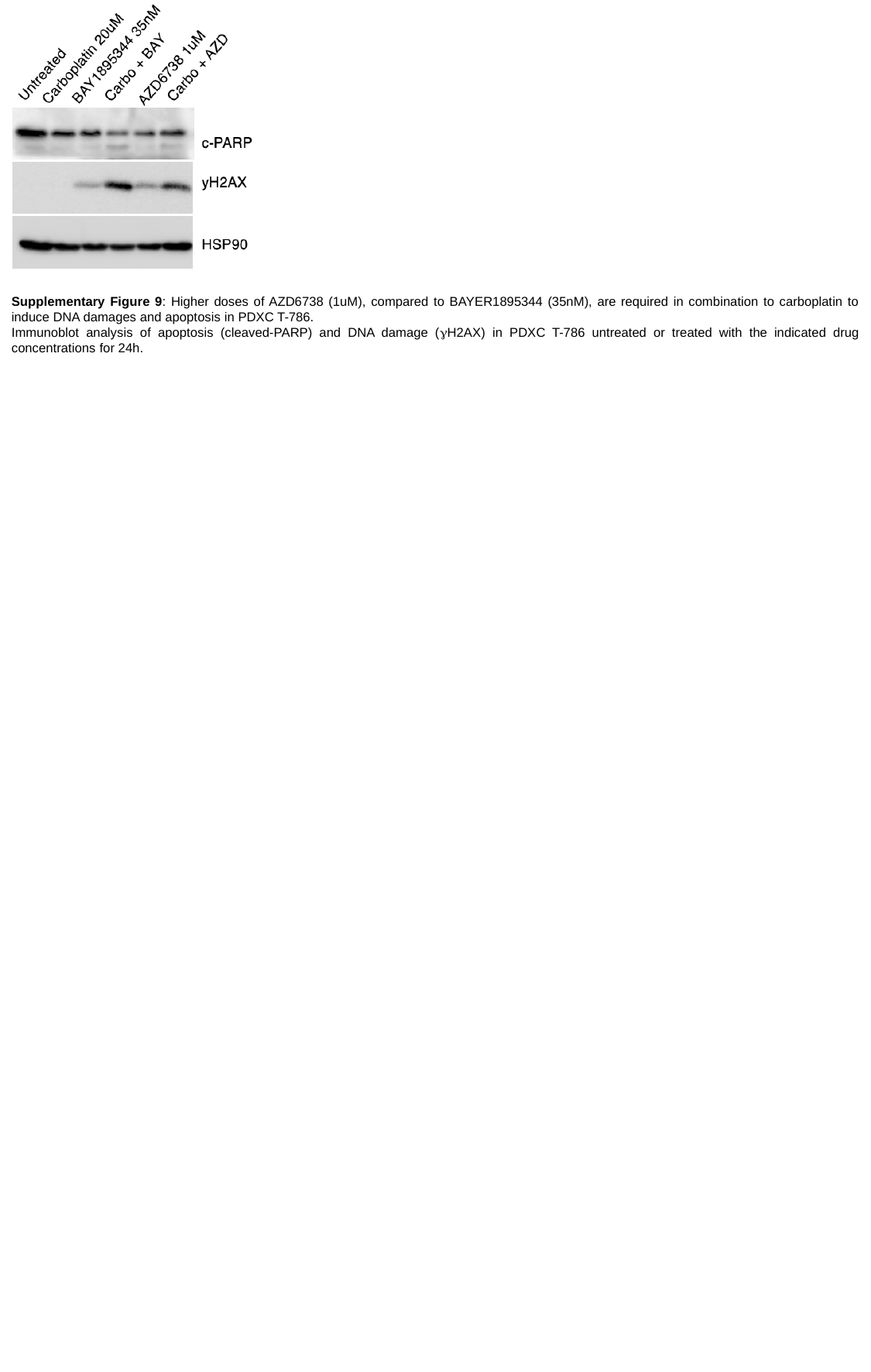

Supplementary Figure 9: Higher doses of AZD6738 (1uM), compared to BAYER1895344 (35nM), are required in combination to carboplatin to induce DNA damages and apoptosis in PDXC T-786.
Immunoblot analysis of apoptosis (cleaved-PARP) and DNA damage (γH2AX) in PDXC T-786 untreated or treated with the indicated drug concentrations for 24h.

### Slide 10
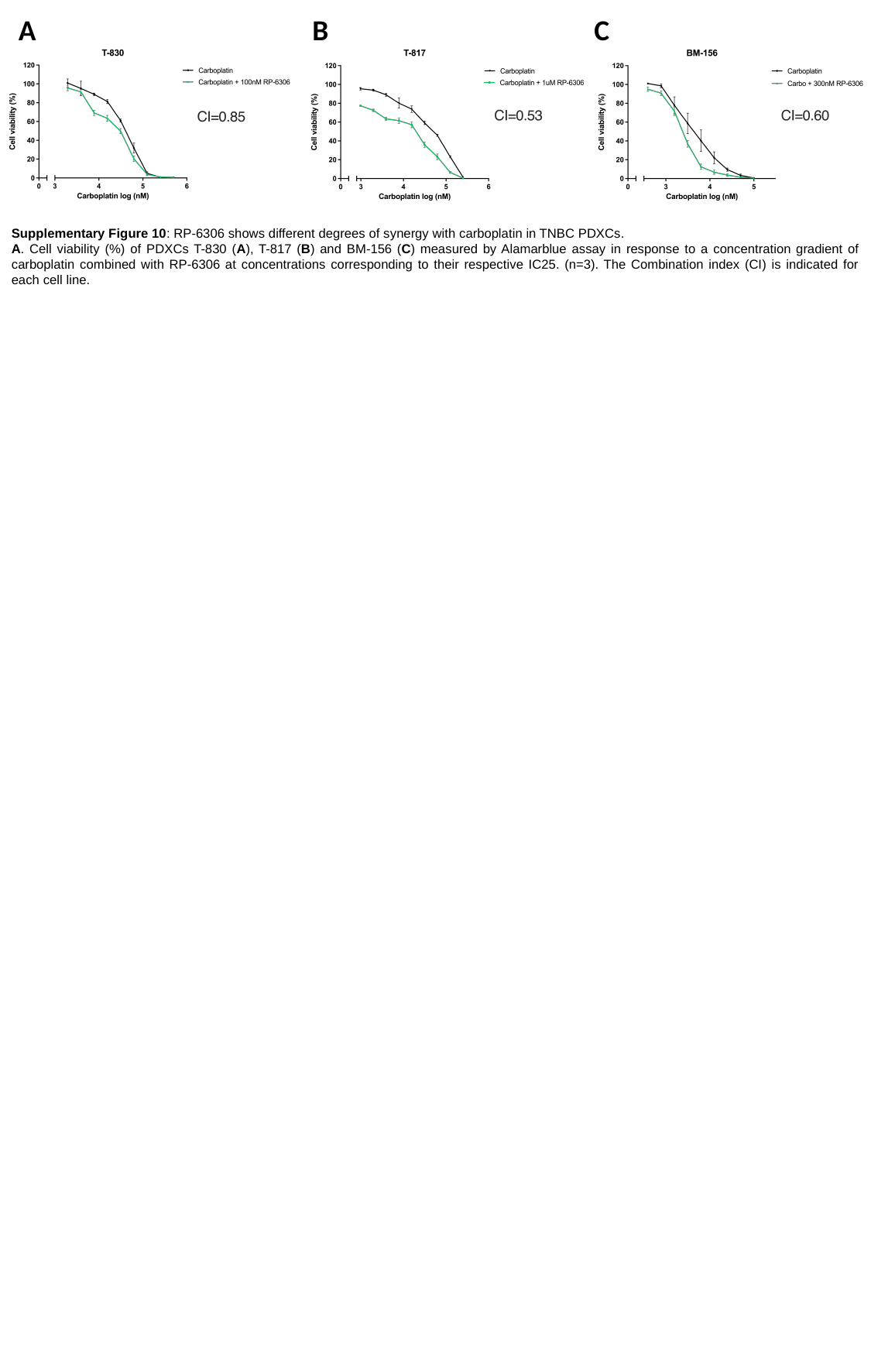

A
B
C
Supplementary Figure 10: RP-6306 shows different degrees of synergy with carboplatin in TNBC PDXCs.
A. Cell viability (%) of PDXCs T-830 (A), T-817 (B) and BM-156 (C) measured by Alamarblue assay in response to a concentration gradient of carboplatin combined with RP-6306 at concentrations corresponding to their respective IC25. (n=3). The Combination index (CI) is indicated for each cell line.

### Slide 11
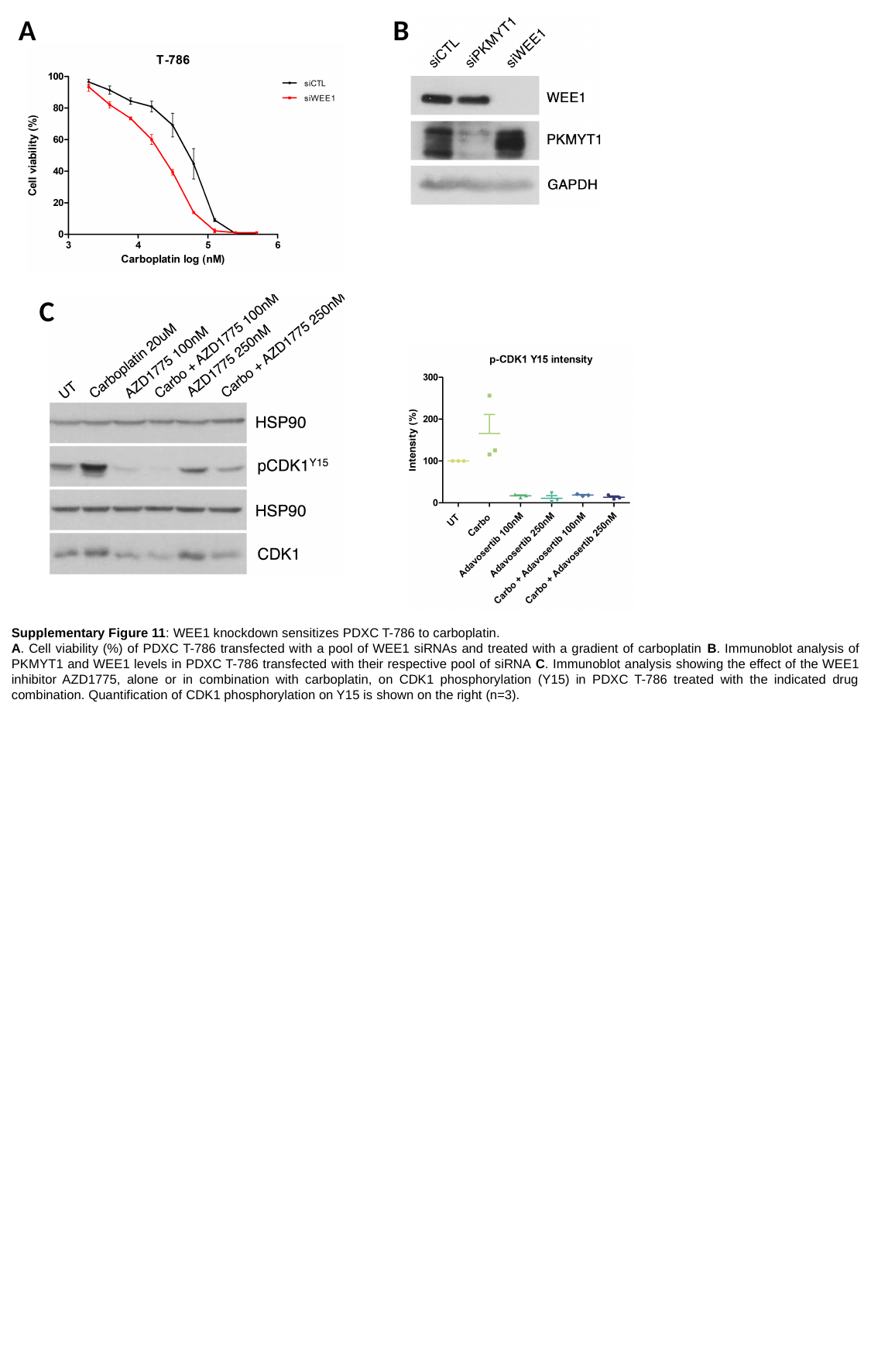

A
B
C
Supplementary Figure 11: WEE1 knockdown sensitizes PDXC T-786 to carboplatin.
A. Cell viability (%) of PDXC T-786 transfected with a pool of WEE1 siRNAs and treated with a gradient of carboplatin B. Immunoblot analysis of PKMYT1 and WEE1 levels in PDXC T-786 transfected with their respective pool of siRNA C. Immunoblot analysis showing the effect of the WEE1 inhibitor AZD1775, alone or in combination with carboplatin, on CDK1 phosphorylation (Y15) in PDXC T-786 treated with the indicated drug combination. Quantification of CDK1 phosphorylation on Y15 is shown on the right (n=3).

### Slide 12
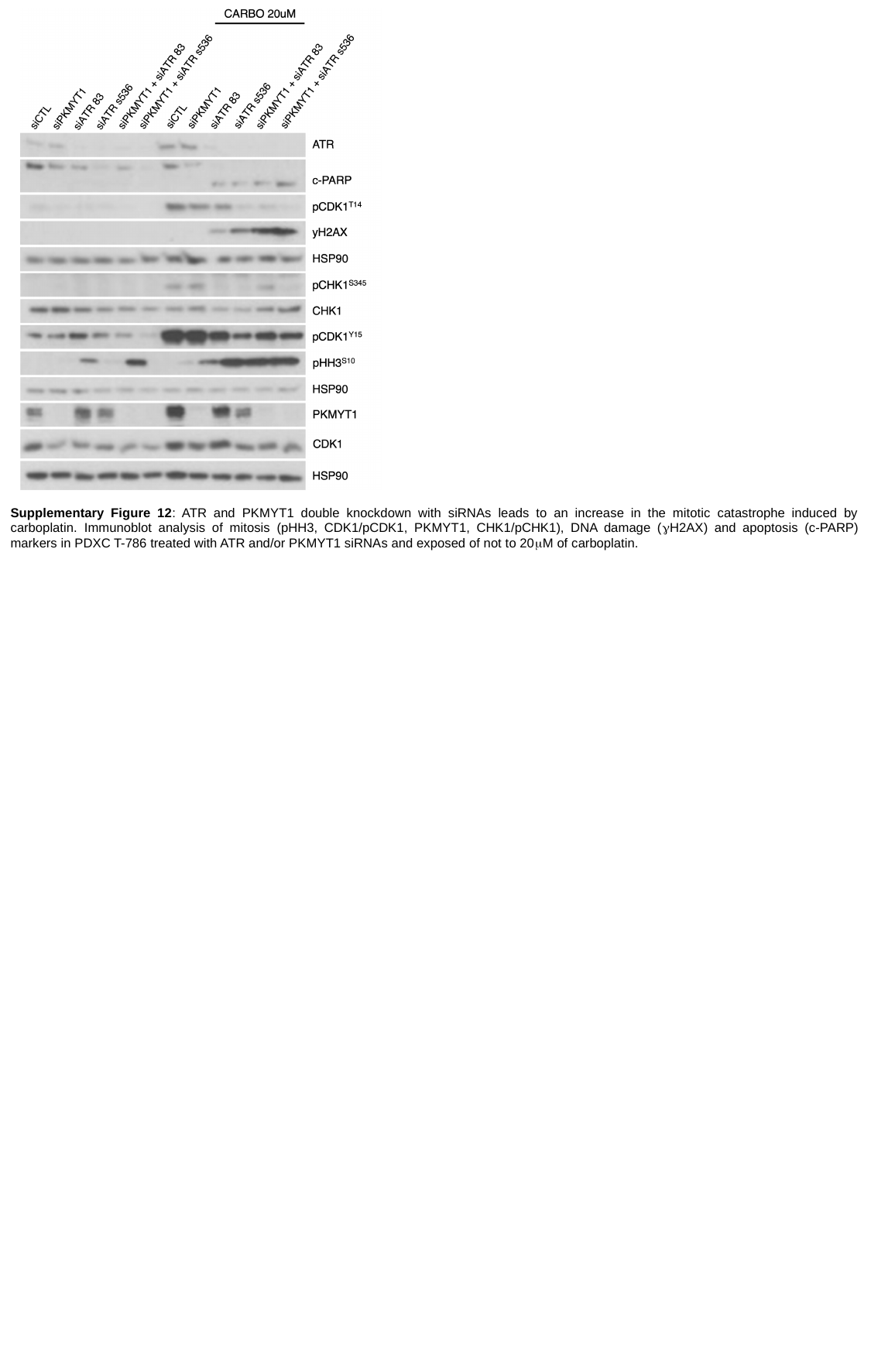

Supplementary Figure 12: ATR and PKMYT1 double knockdown with siRNAs leads to an increase in the mitotic catastrophe induced by carboplatin. Immunoblot analysis of mitosis (pHH3, CDK1/pCDK1, PKMYT1, CHK1/pCHK1), DNA damage (γH2AX) and apoptosis (c-PARP) markers in PDXC T-786 treated with ATR and/or PKMYT1 siRNAs and exposed of not to 20μM of carboplatin.

### Slide 13
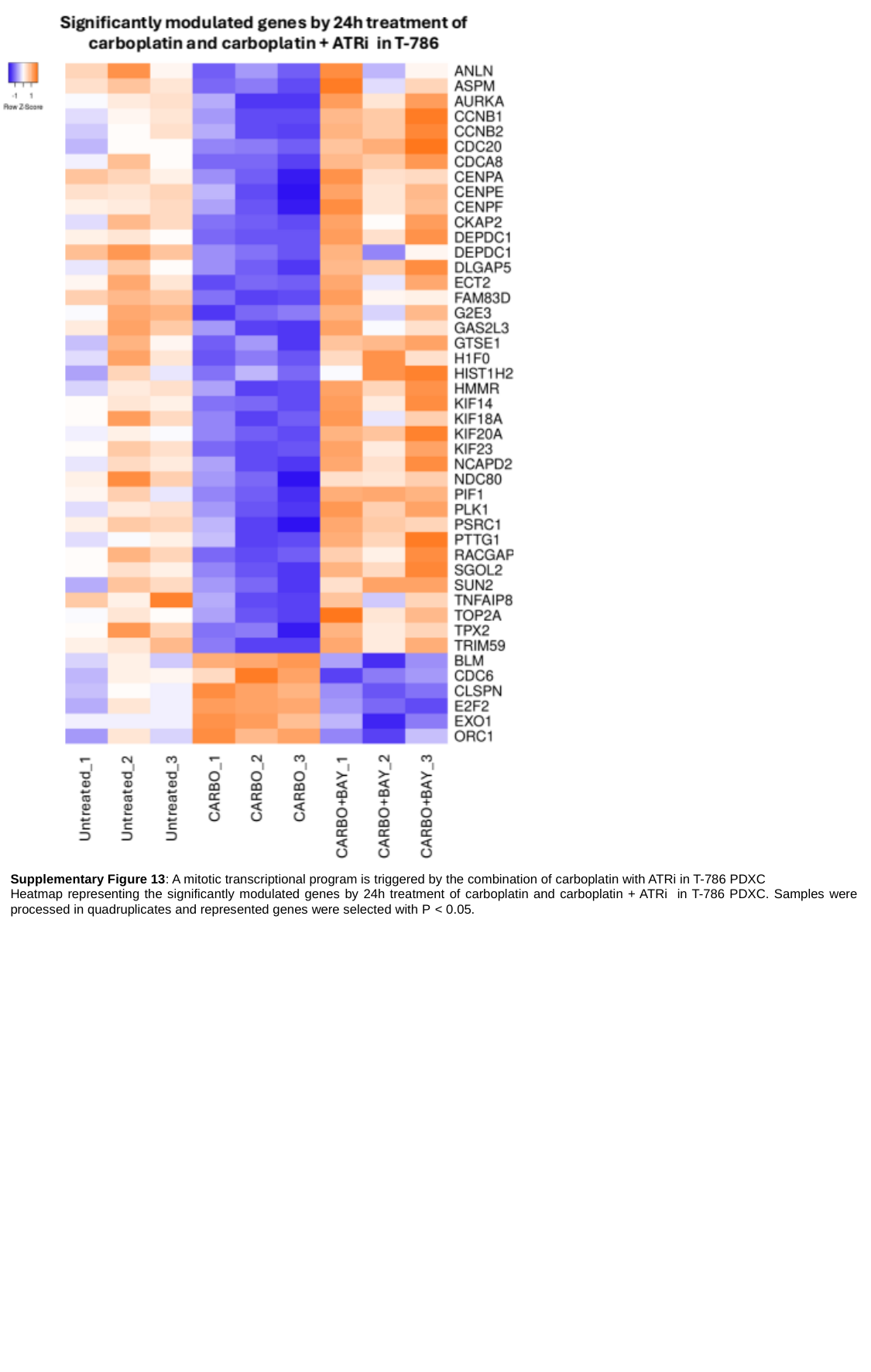

Supplementary Figure 13: A mitotic transcriptional program is triggered by the combination of carboplatin with ATRi in T-786 PDXC
Heatmap representing the significantly modulated genes by 24h treatment of carboplatin and carboplatin + ATRi in T-786 PDXC. Samples were processed in quadruplicates and represented genes were selected with P < 0.05.

### Slide 14
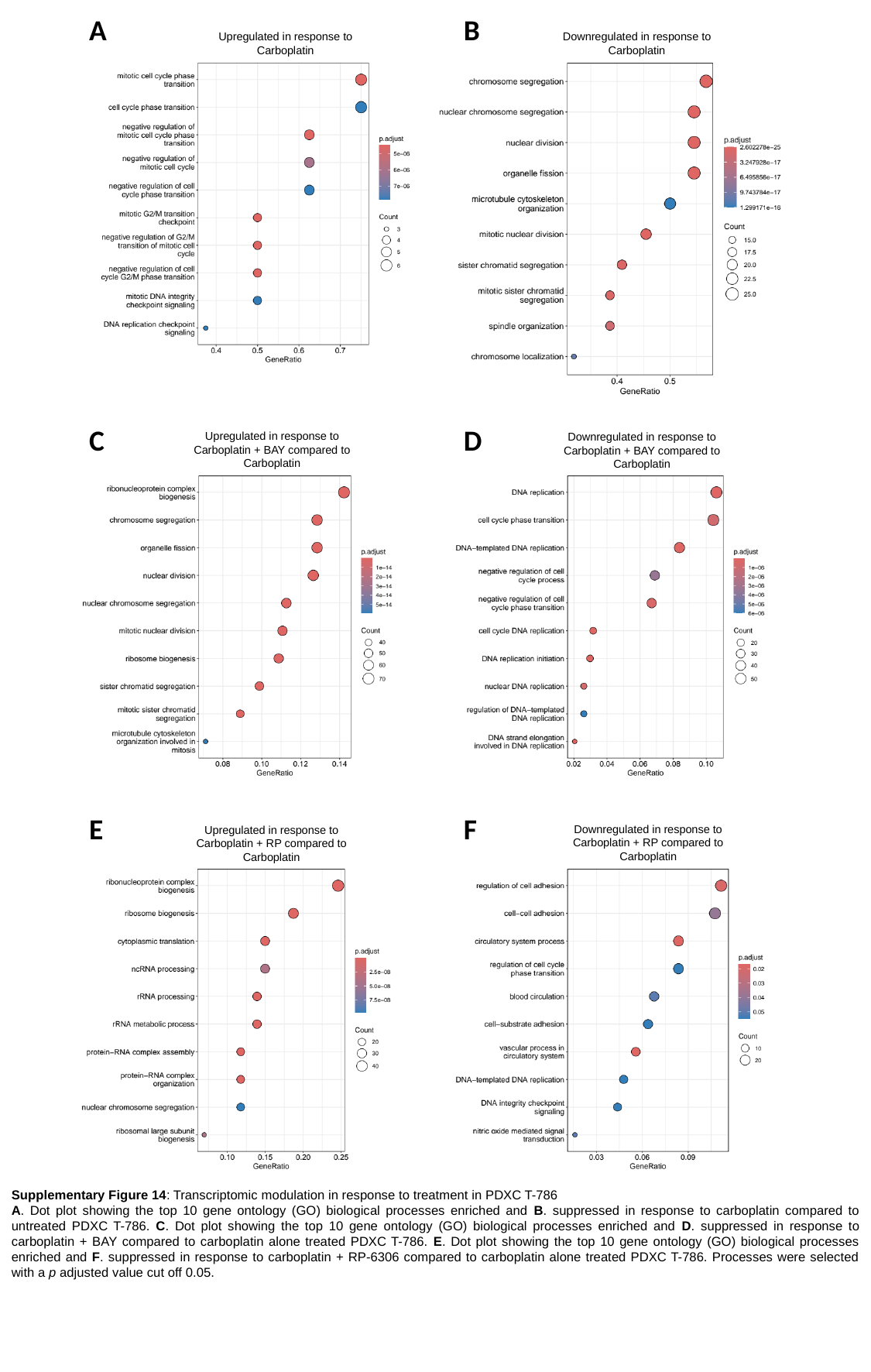

A
B
Upregulated in response to Carboplatin
Downregulated in response to Carboplatin
C
D
Upregulated in response to Carboplatin + BAY compared to Carboplatin
Downregulated in response to Carboplatin + BAY compared to Carboplatin
E
F
Downregulated in response to Carboplatin + RP compared to Carboplatin
Upregulated in response to Carboplatin + RP compared to Carboplatin
Supplementary Figure 14: Transcriptomic modulation in response to treatment in PDXC T-786
A. Dot plot showing the top 10 gene ontology (GO) biological processes enriched and B. suppressed in response to carboplatin compared to untreated PDXC T-786. C. Dot plot showing the top 10 gene ontology (GO) biological processes enriched and D. suppressed in response to carboplatin + BAY compared to carboplatin alone treated PDXC T-786. E. Dot plot showing the top 10 gene ontology (GO) biological processes enriched and F. suppressed in response to carboplatin + RP-6306 compared to carboplatin alone treated PDXC T-786. Processes were selected with a p adjusted value cut off 0.05.

### Slide 15
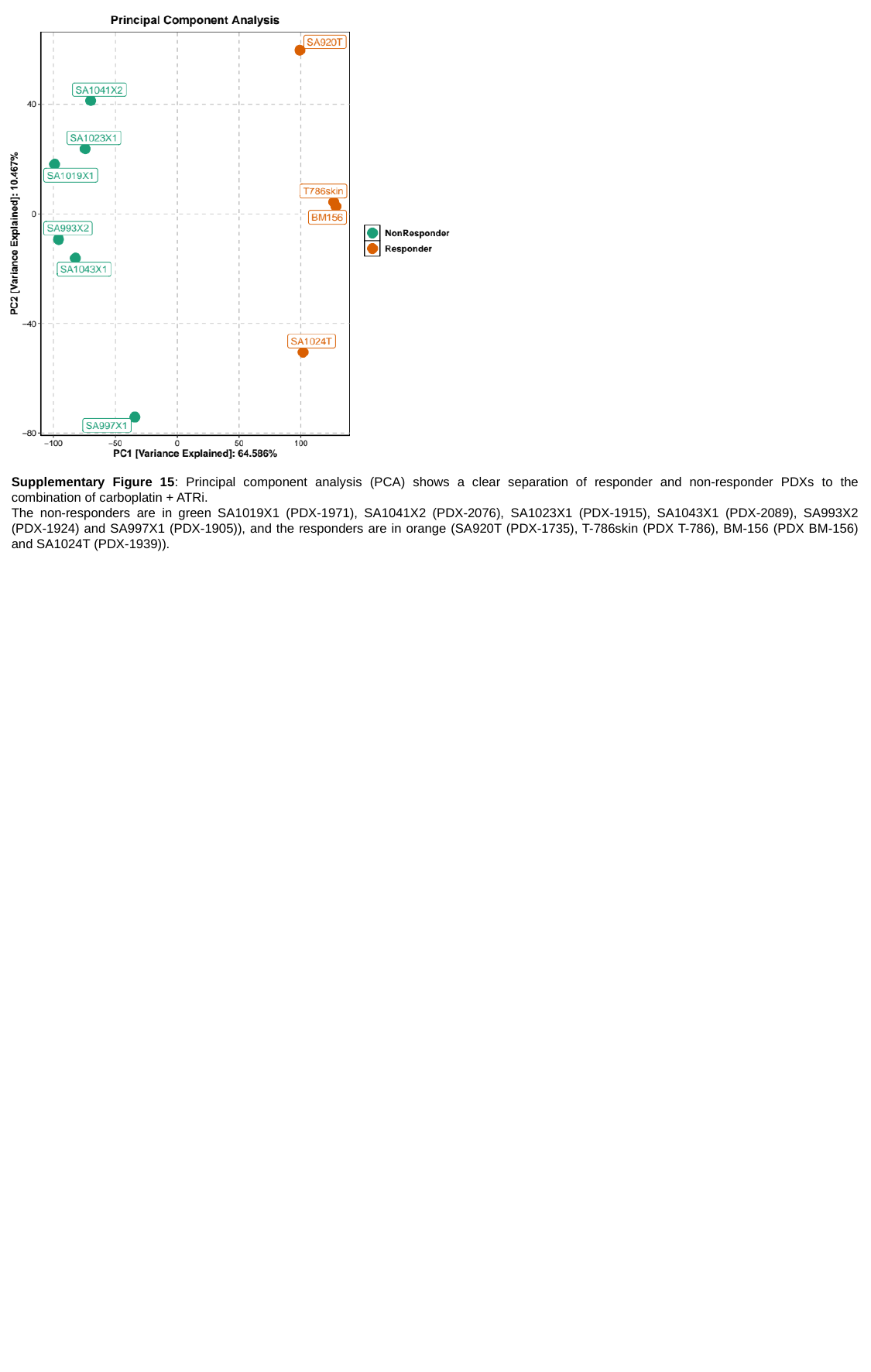

Supplementary Figure 15: Principal component analysis (PCA) shows a clear separation of responder and non-responder PDXs to the combination of carboplatin + ATRi.
The non-responders are in green SA1019X1 (PDX-1971), SA1041X2 (PDX-2076), SA1023X1 (PDX-1915), SA1043X1 (PDX-2089), SA993X2 (PDX-1924) and SA997X1 (PDX-1905)), and the responders are in orange (SA920T (PDX-1735), T-786skin (PDX T-786), BM-156 (PDX BM-156) and SA1024T (PDX-1939)).

### Slide 16
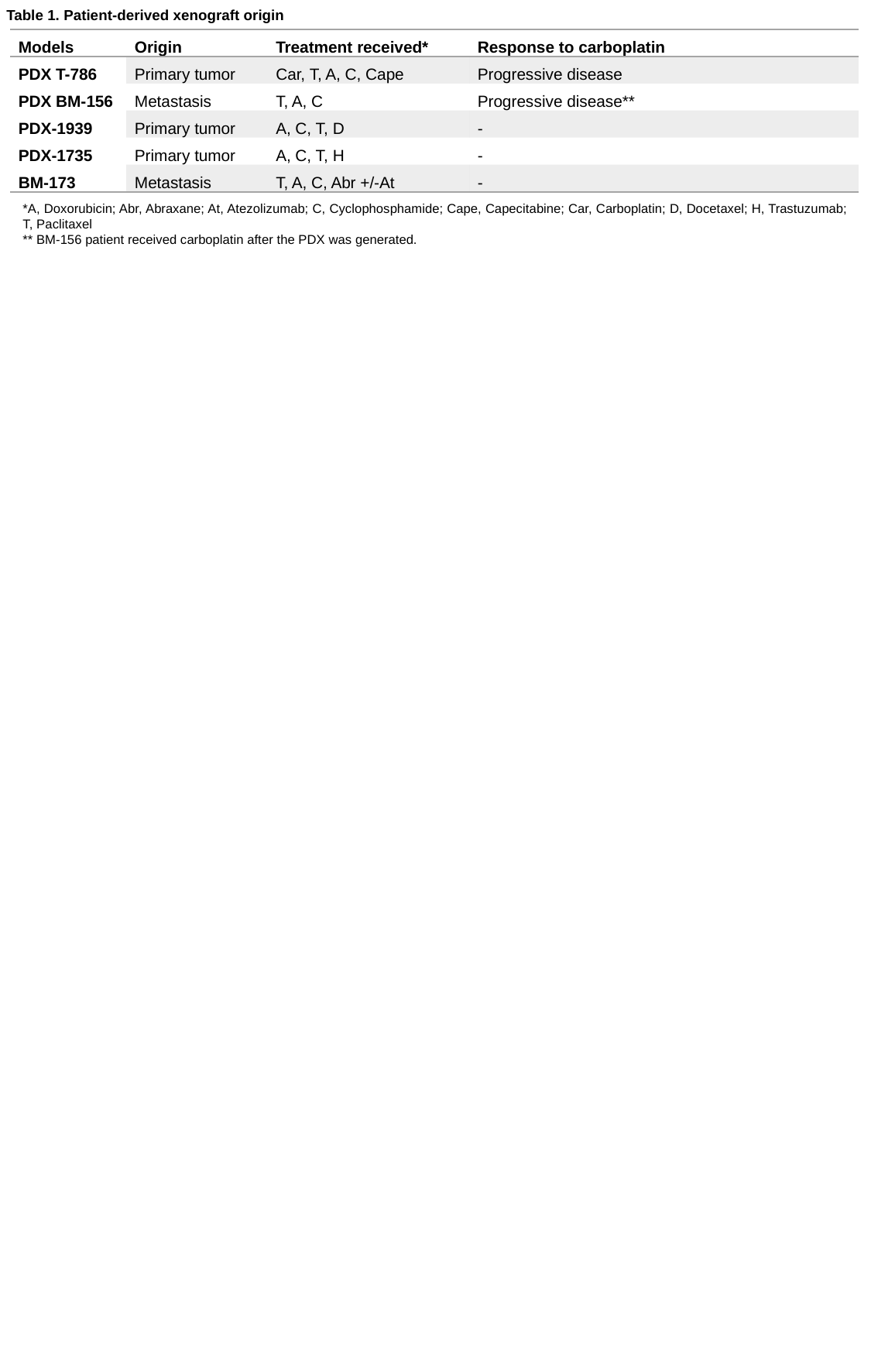

Table 1. Patient-derived xenograft origin
| Models | Origin | Treatment received\* | Response to carboplatin |
| --- | --- | --- | --- |
| PDX T-786 | Primary tumor | Car, T, A, C, Cape | Progressive disease |
| PDX BM-156 | Metastasis | T, A, C | Progressive disease\*\* |
| PDX-1939 | Primary tumor | A, C, T, D | - |
| PDX-1735 | Primary tumor | A, C, T, H | - |
| BM-173 | Metastasis | T, A, C, Abr +/-At | - |
*A, Doxorubicin; Abr, Abraxane; At, Atezolizumab; C, Cyclophosphamide; Cape, Capecitabine; Car, Carboplatin; D, Docetaxel; H, Trastuzumab; T, Paclitaxel
** BM-156 patient received carboplatin after the PDX was generated.

### Slide 17
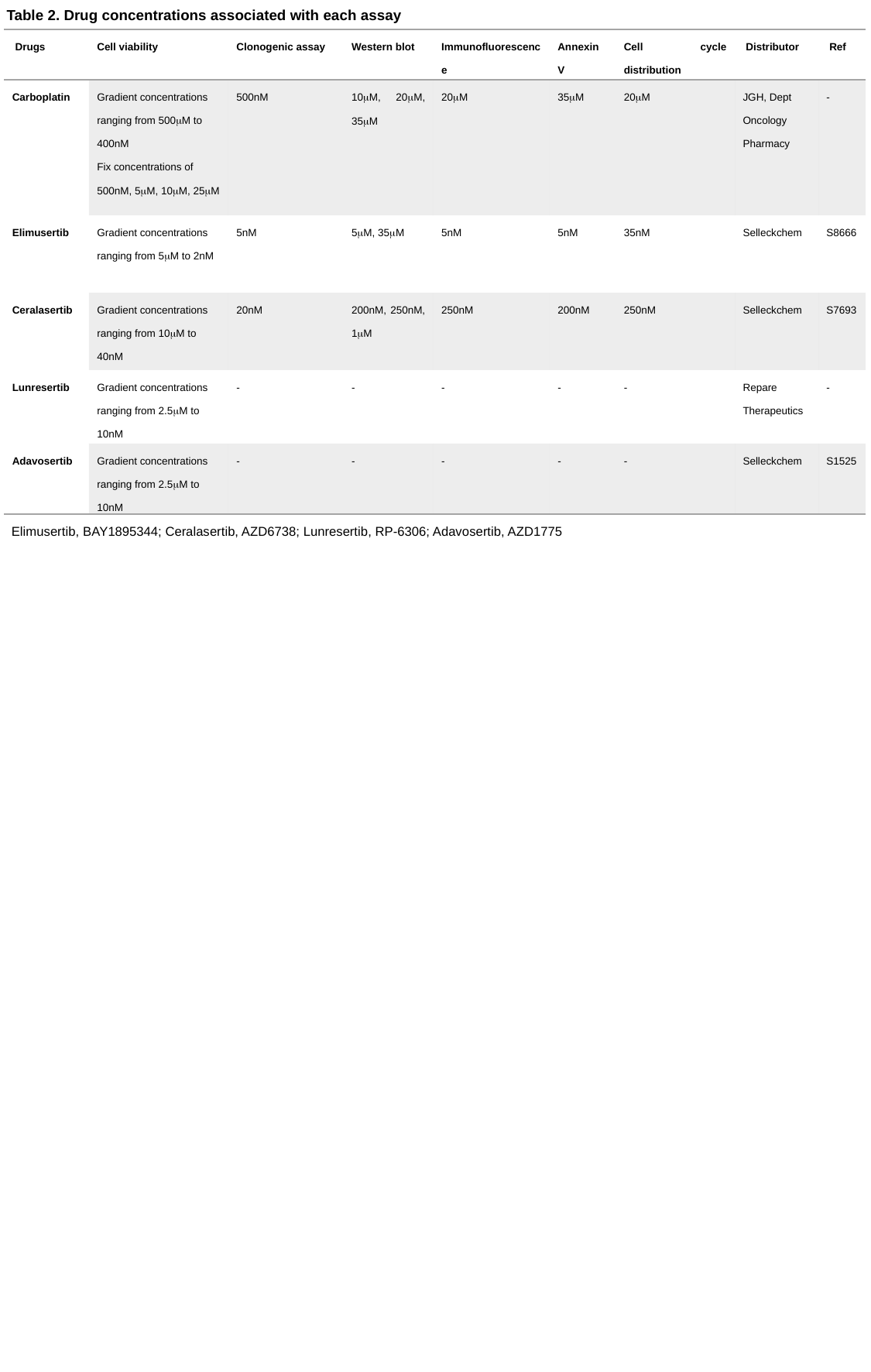

Table 2. Drug concentrations associated with each assay
| Drugs | Cell viability | Clonogenic assay | Western blot | Immunofluorescence | Annexin V | Cell cycle distribution | Distributor | Ref |
| --- | --- | --- | --- | --- | --- | --- | --- | --- |
| Carboplatin | Gradient concentrations ranging from 500μM to 400nM Fix concentrations of 500nM, 5μM, 10μM, 25μM | 500nM | 10μM, 20μM, 35μM | 20μM | 35μM | 20μM | JGH, Dept Oncology Pharmacy | - |
| Elimusertib | Gradient concentrations ranging from 5μM to 2nM | 5nM | 5μM, 35μM | 5nM | 5nM | 35nM | Selleckchem | S8666 |
| Ceralasertib | Gradient concentrations ranging from 10μM to 40nM | 20nM | 200nM, 250nM, 1μM | 250nM | 200nM | 250nM | Selleckchem | S7693 |
| Lunresertib | Gradient concentrations ranging from 2.5μM to 10nM | - | - | - | - | - | Repare Therapeutics | - |
| Adavosertib | Gradient concentrations ranging from 2.5μM to 10nM | - | - | - | - | - | Selleckchem | S1525 |
Elimusertib, BAY1895344; Ceralasertib, AZD6738; Lunresertib, RP-6306; Adavosertib, AZD1775

### Slide 18
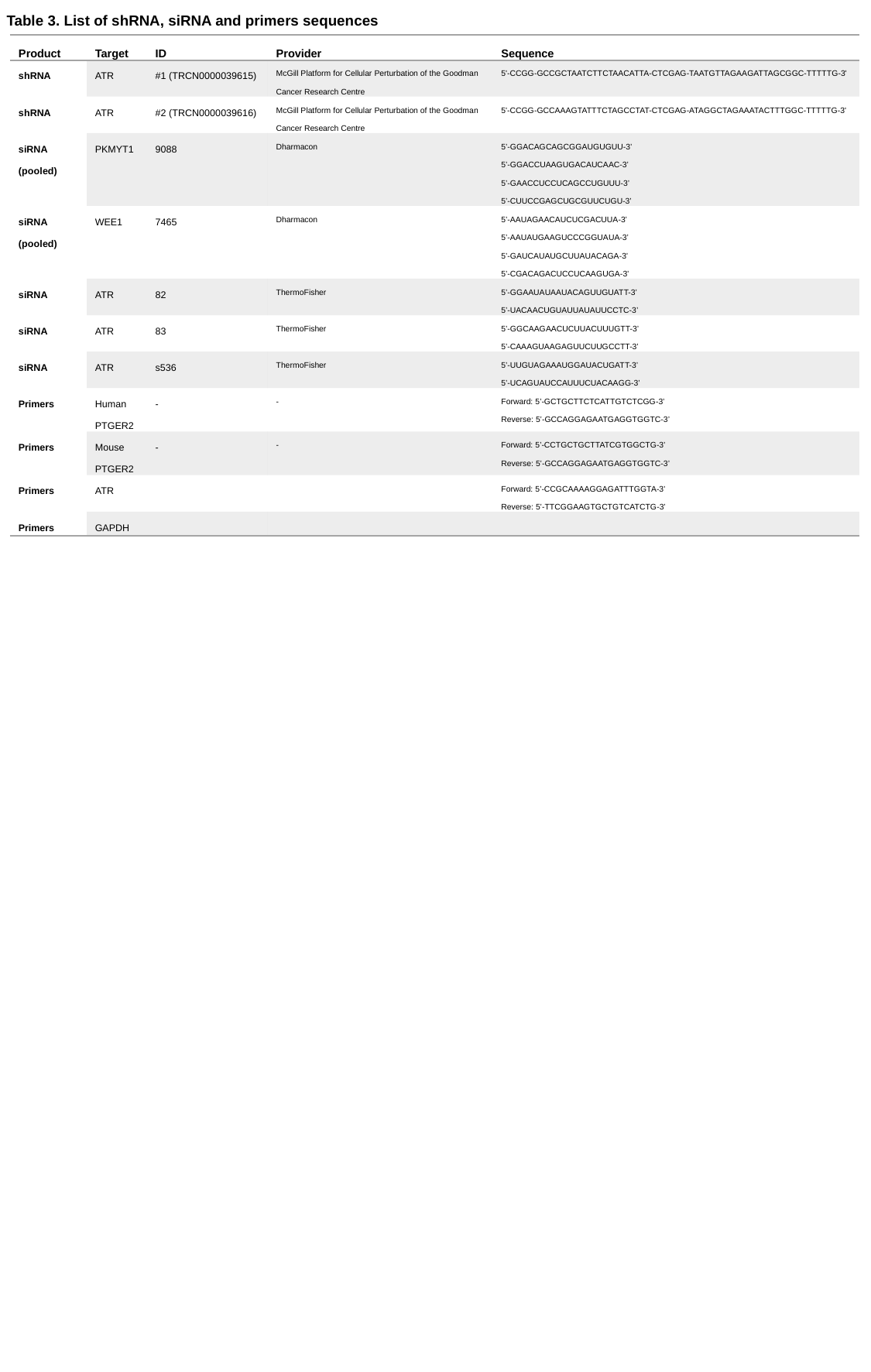

Table 3. List of shRNA, siRNA and primers sequences
| Product | Target | ID | Provider | Sequence |
| --- | --- | --- | --- | --- |
| shRNA | ATR | #1 (TRCN0000039615) | McGill Platform for Cellular Perturbation of the Goodman Cancer Research Centre | 5'-CCGG-GCCGCTAATCTTCTAACATTA-CTCGAG-TAATGTTAGAAGATTAGCGGC-TTTTTG-3' |
| shRNA | ATR | #2 (TRCN0000039616) | McGill Platform for Cellular Perturbation of the Goodman Cancer Research Centre | 5'-CCGG-GCCAAAGTATTTCTAGCCTAT-CTCGAG-ATAGGCTAGAAATACTTTGGC-TTTTTG-3' |
| siRNA (pooled) | PKMYT1 | 9088 | Dharmacon | 5’-GGACAGCAGCGGAUGUGUU-3’ 5’-GGACCUAAGUGACAUCAAC-3’ 5’-GAACCUCCUCAGCCUGUUU-3’ 5’-CUUCCGAGCUGCGUUCUGU-3’ |
| siRNA (pooled) | WEE1 | 7465 | Dharmacon | 5’-AAUAGAACAUCUCGACUUA-3’ 5’-AAUAUGAAGUCCCGGUAUA-3’ 5’-GAUCAUAUGCUUAUACAGA-3’ 5’-CGACAGACUCCUCAAGUGA-3’ |
| siRNA | ATR | 82 | ThermoFisher | 5’-GGAAUAUAAUACAGUUGUATT-3’ 5’-UACAACUGUAUUAUAUUCCTC-3’ |
| siRNA | ATR | 83 | ThermoFisher | 5’-GGCAAGAACUCUUACUUUGTT-3’ 5’-CAAAGUAAGAGUUCUUGCCTT-3’ |
| siRNA | ATR | s536 | ThermoFisher | 5’-UUGUAGAAAUGGAUACUGATT-3’ 5’-UCAGUAUCCAUUUCUACAAGG-3’ |
| Primers | Human PTGER2 | - | - | Forward: 5’-GCTGCTTCTCATTGTCTCGG-3’ Reverse: 5’-GCCAGGAGAATGAGGTGGTC-3’ |
| Primers | Mouse PTGER2 | - | - | Forward: 5’-CCTGCTGCTTATCGTGGCTG-3’ Reverse: 5’-GCCAGGAGAATGAGGTGGTC-3’ |
| Primers | ATR | | | Forward: 5’-CCGCAAAAGGAGATTTGGTA-3’ Reverse: 5’-TTCGGAAGTGCTGTCATCTG-3’ |
| Primers | GAPDH | | | |

### Slide 19
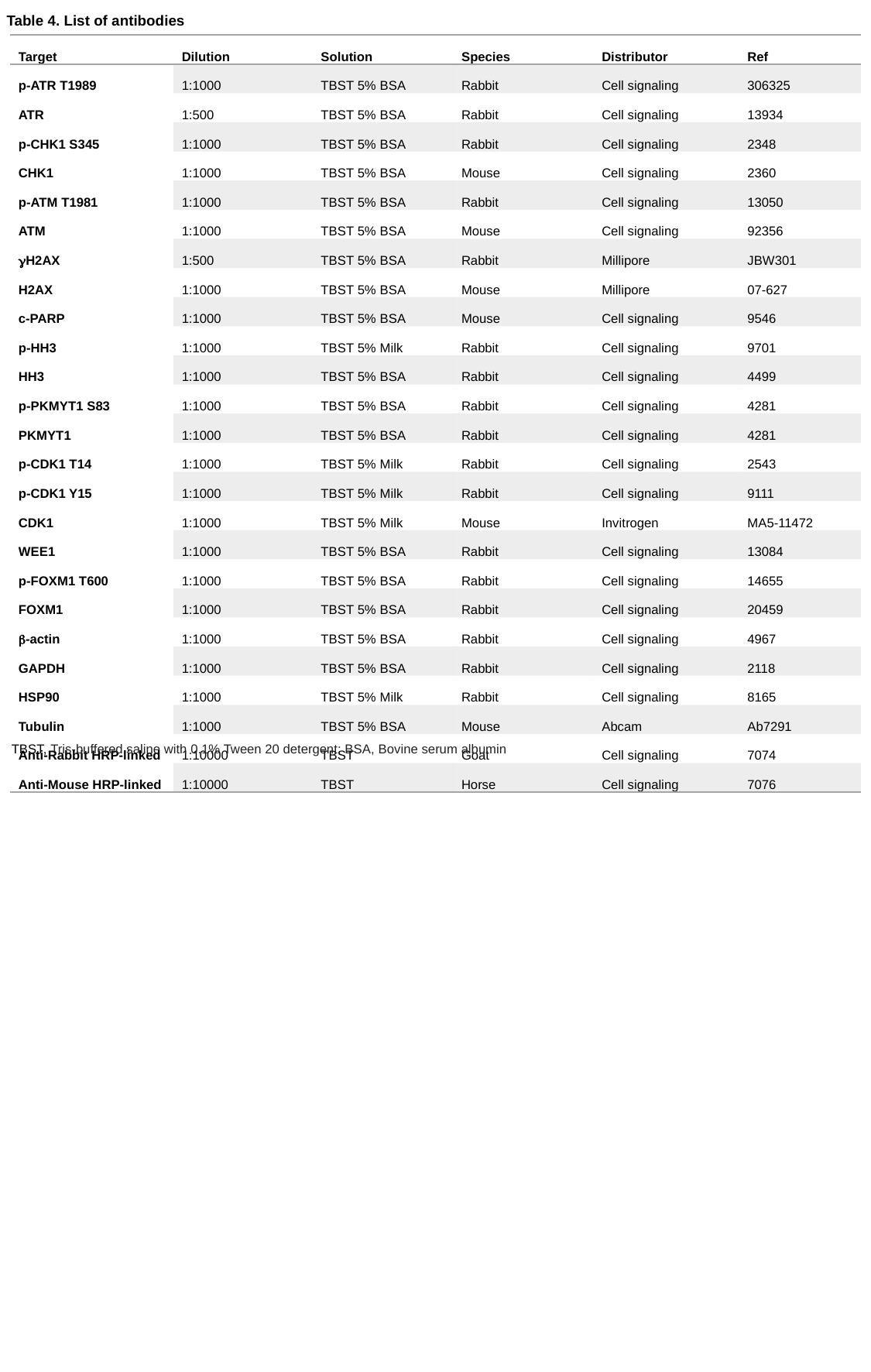

Table 4. List of antibodies
| Target | Dilution | Solution | Species | Distributor | Ref |
| --- | --- | --- | --- | --- | --- |
| p-ATR T1989 | 1:1000 | TBST 5% BSA | Rabbit | Cell signaling | 306325 |
| ATR | 1:500 | TBST 5% BSA | Rabbit | Cell signaling | 13934 |
| p-CHK1 S345 | 1:1000 | TBST 5% BSA | Rabbit | Cell signaling | 2348 |
| CHK1 | 1:1000 | TBST 5% BSA | Mouse | Cell signaling | 2360 |
| p-ATM T1981 | 1:1000 | TBST 5% BSA | Rabbit | Cell signaling | 13050 |
| ATM | 1:1000 | TBST 5% BSA | Mouse | Cell signaling | 92356 |
| γH2AX | 1:500 | TBST 5% BSA | Rabbit | Millipore | JBW301 |
| H2AX | 1:1000 | TBST 5% BSA | Mouse | Millipore | 07-627 |
| c-PARP | 1:1000 | TBST 5% BSA | Mouse | Cell signaling | 9546 |
| p-HH3 | 1:1000 | TBST 5% Milk | Rabbit | Cell signaling | 9701 |
| HH3 | 1:1000 | TBST 5% BSA | Rabbit | Cell signaling | 4499 |
| p-PKMYT1 S83 | 1:1000 | TBST 5% BSA | Rabbit | Cell signaling | 4281 |
| PKMYT1 | 1:1000 | TBST 5% BSA | Rabbit | Cell signaling | 4281 |
| p-CDK1 T14 | 1:1000 | TBST 5% Milk | Rabbit | Cell signaling | 2543 |
| p-CDK1 Y15 | 1:1000 | TBST 5% Milk | Rabbit | Cell signaling | 9111 |
| CDK1 | 1:1000 | TBST 5% Milk | Mouse | Invitrogen | MA5-11472 |
| WEE1 | 1:1000 | TBST 5% BSA | Rabbit | Cell signaling | 13084 |
| p-FOXM1 T600 | 1:1000 | TBST 5% BSA | Rabbit | Cell signaling | 14655 |
| FOXM1 | 1:1000 | TBST 5% BSA | Rabbit | Cell signaling | 20459 |
| β-actin | 1:1000 | TBST 5% BSA | Rabbit | Cell signaling | 4967 |
| GAPDH | 1:1000 | TBST 5% BSA | Rabbit | Cell signaling | 2118 |
| HSP90 | 1:1000 | TBST 5% Milk | Rabbit | Cell signaling | 8165 |
| Tubulin | 1:1000 | TBST 5% BSA | Mouse | Abcam | Ab7291 |
| Anti-Rabbit HRP-linked | 1:10000 | TBST | Goat | Cell signaling | 7074 |
| Anti-Mouse HRP-linked | 1:10000 | TBST | Horse | Cell signaling | 7076 |
TBST, Tris-buffered saline with 0.1% Tween 20 detergent; BSA, Bovine serum albumin

### Slide 20
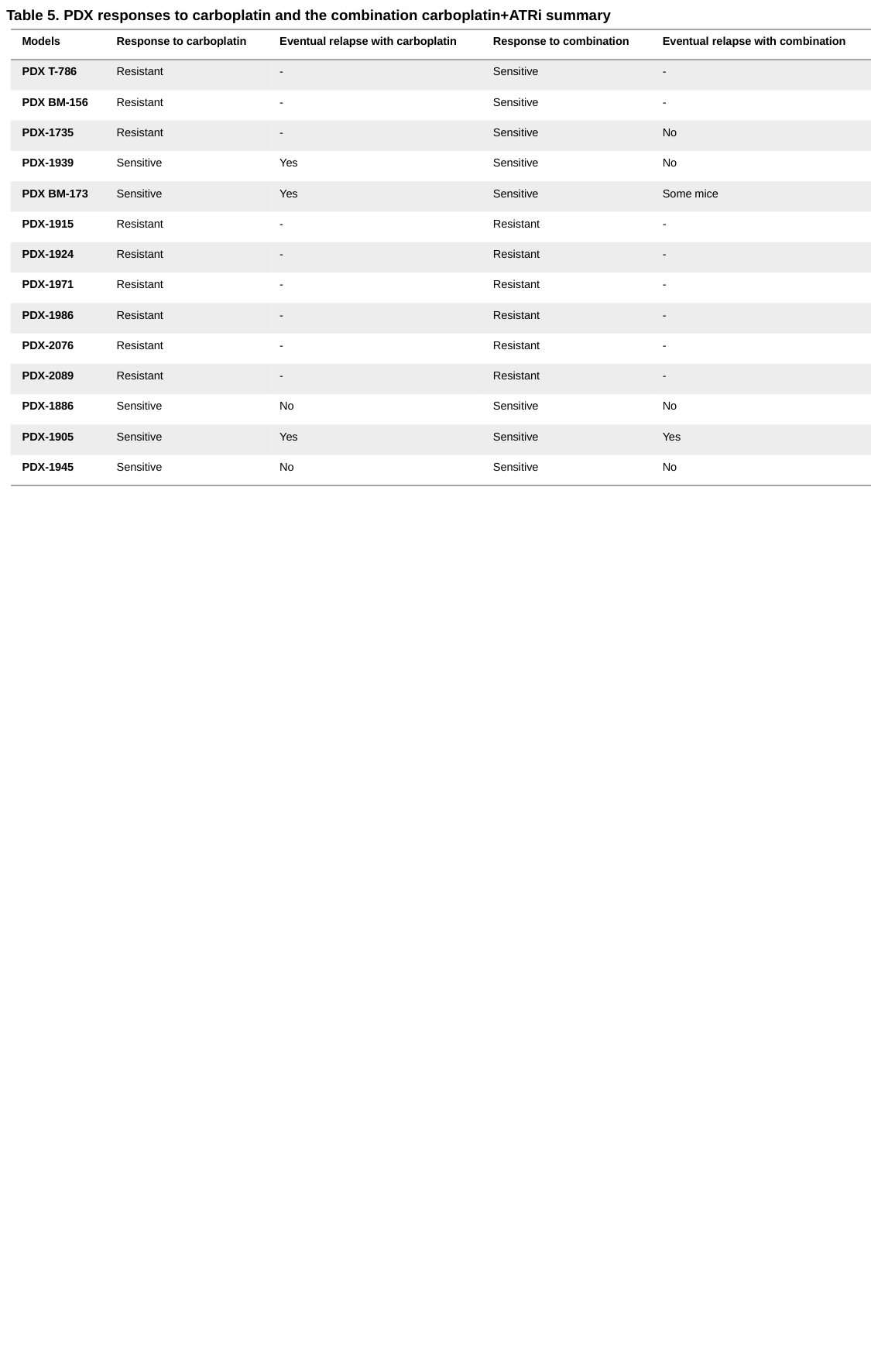

Table 5. PDX responses to carboplatin and the combination carboplatin+ATRi summary
| Models | Response to carboplatin | Eventual relapse with carboplatin | Response to combination | Eventual relapse with combination |
| --- | --- | --- | --- | --- |
| PDX T-786 | Resistant | - | Sensitive | - |
| PDX BM-156 | Resistant | - | Sensitive | - |
| PDX-1735 | Resistant | - | Sensitive | No |
| PDX-1939 | Sensitive | Yes | Sensitive | No |
| PDX BM-173 | Sensitive | Yes | Sensitive | Some mice |
| PDX-1915 | Resistant | - | Resistant | - |
| PDX-1924 | Resistant | - | Resistant | - |
| PDX-1971 | Resistant | - | Resistant | - |
| PDX-1986 | Resistant | - | Resistant | - |
| PDX-2076 | Resistant | - | Resistant | - |
| PDX-2089 | Resistant | - | Resistant | - |
| PDX-1886 | Sensitive | No | Sensitive | No |
| PDX-1905 | Sensitive | Yes | Sensitive | Yes |
| PDX-1945 | Sensitive | No | Sensitive | No |

### Slide 21
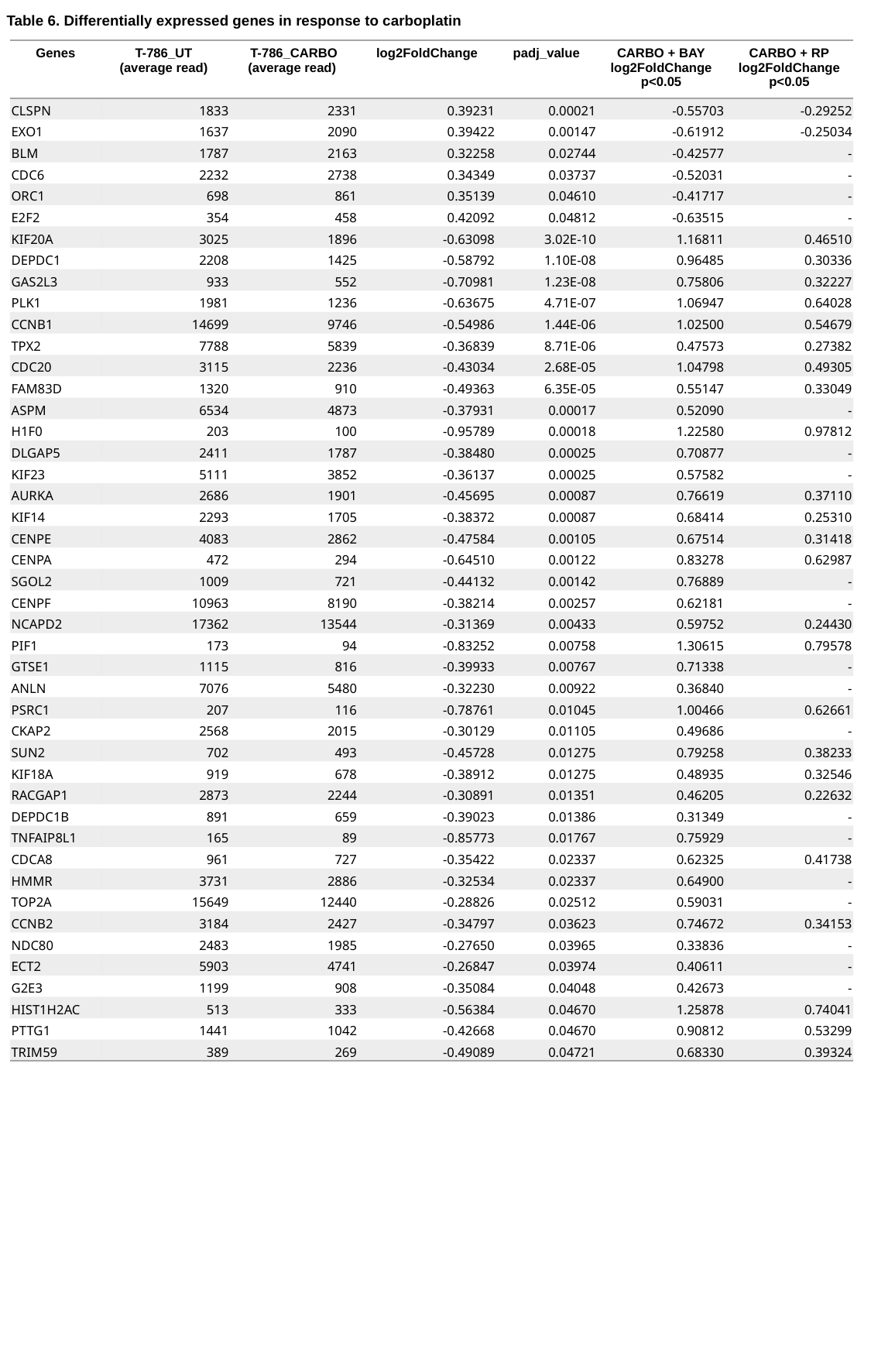

Table 6. Differentially expressed genes in response to carboplatin
| Genes | T-786\_UT (average read) | T-786\_CARBO (average read) | log2FoldChange | padj\_value | CARBO + BAY log2FoldChange p<0.05 | CARBO + RP log2FoldChange p<0.05 |
| --- | --- | --- | --- | --- | --- | --- |
| CLSPN | 1833 | 2331 | 0.39231 | 0.00021 | -0.55703 | -0.29252 |
| EXO1 | 1637 | 2090 | 0.39422 | 0.00147 | -0.61912 | -0.25034 |
| BLM | 1787 | 2163 | 0.32258 | 0.02744 | -0.42577 | - |
| CDC6 | 2232 | 2738 | 0.34349 | 0.03737 | -0.52031 | - |
| ORC1 | 698 | 861 | 0.35139 | 0.04610 | -0.41717 | - |
| E2F2 | 354 | 458 | 0.42092 | 0.04812 | -0.63515 | - |
| KIF20A | 3025 | 1896 | -0.63098 | 3.02E-10 | 1.16811 | 0.46510 |
| DEPDC1 | 2208 | 1425 | -0.58792 | 1.10E-08 | 0.96485 | 0.30336 |
| GAS2L3 | 933 | 552 | -0.70981 | 1.23E-08 | 0.75806 | 0.32227 |
| PLK1 | 1981 | 1236 | -0.63675 | 4.71E-07 | 1.06947 | 0.64028 |
| CCNB1 | 14699 | 9746 | -0.54986 | 1.44E-06 | 1.02500 | 0.54679 |
| TPX2 | 7788 | 5839 | -0.36839 | 8.71E-06 | 0.47573 | 0.27382 |
| CDC20 | 3115 | 2236 | -0.43034 | 2.68E-05 | 1.04798 | 0.49305 |
| FAM83D | 1320 | 910 | -0.49363 | 6.35E-05 | 0.55147 | 0.33049 |
| ASPM | 6534 | 4873 | -0.37931 | 0.00017 | 0.52090 | - |
| H1F0 | 203 | 100 | -0.95789 | 0.00018 | 1.22580 | 0.97812 |
| DLGAP5 | 2411 | 1787 | -0.38480 | 0.00025 | 0.70877 | - |
| KIF23 | 5111 | 3852 | -0.36137 | 0.00025 | 0.57582 | - |
| AURKA | 2686 | 1901 | -0.45695 | 0.00087 | 0.76619 | 0.37110 |
| KIF14 | 2293 | 1705 | -0.38372 | 0.00087 | 0.68414 | 0.25310 |
| CENPE | 4083 | 2862 | -0.47584 | 0.00105 | 0.67514 | 0.31418 |
| CENPA | 472 | 294 | -0.64510 | 0.00122 | 0.83278 | 0.62987 |
| SGOL2 | 1009 | 721 | -0.44132 | 0.00142 | 0.76889 | - |
| CENPF | 10963 | 8190 | -0.38214 | 0.00257 | 0.62181 | - |
| NCAPD2 | 17362 | 13544 | -0.31369 | 0.00433 | 0.59752 | 0.24430 |
| PIF1 | 173 | 94 | -0.83252 | 0.00758 | 1.30615 | 0.79578 |
| GTSE1 | 1115 | 816 | -0.39933 | 0.00767 | 0.71338 | - |
| ANLN | 7076 | 5480 | -0.32230 | 0.00922 | 0.36840 | - |
| PSRC1 | 207 | 116 | -0.78761 | 0.01045 | 1.00466 | 0.62661 |
| CKAP2 | 2568 | 2015 | -0.30129 | 0.01105 | 0.49686 | - |
| SUN2 | 702 | 493 | -0.45728 | 0.01275 | 0.79258 | 0.38233 |
| KIF18A | 919 | 678 | -0.38912 | 0.01275 | 0.48935 | 0.32546 |
| RACGAP1 | 2873 | 2244 | -0.30891 | 0.01351 | 0.46205 | 0.22632 |
| DEPDC1B | 891 | 659 | -0.39023 | 0.01386 | 0.31349 | - |
| TNFAIP8L1 | 165 | 89 | -0.85773 | 0.01767 | 0.75929 | - |
| CDCA8 | 961 | 727 | -0.35422 | 0.02337 | 0.62325 | 0.41738 |
| HMMR | 3731 | 2886 | -0.32534 | 0.02337 | 0.64900 | - |
| TOP2A | 15649 | 12440 | -0.28826 | 0.02512 | 0.59031 | - |
| CCNB2 | 3184 | 2427 | -0.34797 | 0.03623 | 0.74672 | 0.34153 |
| NDC80 | 2483 | 1985 | -0.27650 | 0.03965 | 0.33836 | - |
| ECT2 | 5903 | 4741 | -0.26847 | 0.03974 | 0.40611 | - |
| G2E3 | 1199 | 908 | -0.35084 | 0.04048 | 0.42673 | - |
| HIST1H2AC | 513 | 333 | -0.56384 | 0.04670 | 1.25878 | 0.74041 |
| PTTG1 | 1441 | 1042 | -0.42668 | 0.04670 | 0.90812 | 0.53299 |
| TRIM59 | 389 | 269 | -0.49089 | 0.04721 | 0.68330 | 0.39324 |
